# A gatekeeping function of the replicative polymerase controls pathway choice in the resolution of lesion-stalled replisomes

**DOI:** 10.1101/676486

**Authors:** Seungwoo Chang, Karel Naiman, Elizabeth S. Thrall, James E. Kath, Slobodan Jergic, Nicholas Dixon, Robert P. Fuchs, Joseph J Loparo

## Abstract

DNA lesions stall the replisome and proper resolution of these obstructions is critical for genome stability. Replisomes can directly replicate past a lesion by error-prone translesion synthesis. Alternatively, replisomes can reprime DNA synthesis downstream of the lesion, creating a single-stranded DNA gap that is repaired primarily in an error-free, homology-directed manner. Here we demonstrate how structural changes within the bacterial replisome determine the resolution pathway of lesion-stalled replisomes. This pathway selection is controlled by a dynamic interaction between the proofreading subunit of the replicative polymerase and the processivity clamp, which sets a kinetic barrier to restrict access of TLS polymerases to the primer/template junction. Failure of TLS polymerases to overcome this barrier leads to repriming, which competes kinetically with TLS. Our results demonstrate that independent of its exonuclease activity, the proofreading subunit of the replisome acts as a gatekeeper and influences replication fidelity during the resolution of lesion-stalled replisomes.

## Introduction

Genomic DNA is constantly damaged by various intracellular and extracellular agents. Replication is transiently blocked at these sites because replicative DNA polymerases are generally poor at synthesizing past lesions. Stalled replisomes are resolved primarily through either recombination-dependent damage avoidance (DA) pathways or translesion synthesis (TLS), and distinct DNA intermediates are created during each process (Fujii et al., 2018). In DA pathways, damaged templates are replicated in an error-free manner using an undamaged homologous sister chromatid as a template via processes that utilize homologous recombination factors. In contrast, during TLS damaged templates are directly replicated by TLS polymerases in an error-prone manner. TLS can occur through sequential polymerase switching between the replicative polymerase and a TLS polymerase, yielding a continuous DNA product (TLS at the fork). Alternatively, replisomes can reprime DNA synthesis downstream of the lesion leaving a ssDNA gap behind. These gaps are then filled in either by TLS polymerases (Gap-filling synthesis) or by Homology-Dependent Gap Repair (HDGR) (Fujii et al., 2018). Given the marked differences in mutagenic potential between DA pathways and TLS, it is important to understand what determines pathway choice at stalled replisomes.

Structural changes within the replisome upon lesion stalling, such as conformational changes of replisome components and alterations in protein-protein interactions, likely play an important role in pathway selection, yet these dynamics have been largely unexplored. DNA replication of *E. coli* serves as an attractive model system to probe these lesion-induced structural changes of the replisome as it uses both DA pathways and TLS to resolve stalled replisomes. Moreover, the *E. coli* replisome can be reconstituted with a relatively small number of factors and is genetically tractable while retaining the same basic architecture of more complicated systems (Wu et al., 1992a; Yao and O’Donnell, 2010; Zechner et al., 1992a). In *E. coli* the majority of lesion-stalled replisomes are resolved through DA pathways, particularly when the SOS damage response is not induced. However, upon induction of the SOS response and the concomitant increase in TLS polymerase levels, higher fractions of stalled replisomes are resolved through TLS (Fuchs, 2016; Napolitano et al., 2000).

*E. coli* cells have three TLS polymerases, Pol II, IV and V (Napolitano et al., 2000; Sutton, 2010). Among these, Pol II and IV are abundant even before their expression levels are highly elevated during the damage-induced SOS response. If TLS polymerases gained frequent access to the extending primer, replication would be severely inhibited due to the much slower polymerization of TLS polymerases as compared to Pol III (Indiani et al., 2009; Kath et al., 2016; 2014). Intriguingly, despite their high abundance, TLS polymerases only modestly inhibit replication in *E. coli* cells (Tan et al., 2015) and they contribute little to spontaneous mutagenesis (Kuban et al., 2004; McKenzie et al., 2003). Collectively, these observations suggest that TLS polymerases are largely excluded from replisomes (Henrikus et al., 2018; Thrall et al., 2017).

The *E. coli* β_2_ clamp processivity factor plays an important role in regulating TLS as all TLS polymerases must bind the β_2_ clamp to perform TLS (Becherel et al., 2002; Heltzel et al., 2009; Kath et al., 2014; Lenne-Samuel et al., 2002; Napolitano et al., 2000). The β_2_ clamp is a homodimer that encircles DNA and tethers DNA polymerases to their template (Kong et al., 1992; Kuriyan and O’Donnell, 1993). Each protomer of the β_2_ clamp molecule has a hydrophobic cleft, a common binding site for clamp binding proteins, yielding two identical clefts per β_2_ clamp molecule (Bunting et al., 2003). Clamp binding proteins have one or more clamp binding motifs (CBMs), that interact with the β_2_ clamp via cleft-CBM interactions (Dalrymple et al., 2001; Patoli et al., 2013). All five *E. coli* DNA polymerases have one or two conserved CBMs (Becherel et al., 2002; Lamers et al., 2006; Patoli et al., 2013). Pol III, the replicative polymerase, is a trimeric complex (αεθ) consisting of α polymerase, ε exonuclease and θ accessory subunits. The α subunit has an internal CBM, which is required for processive replication (Dohrmann and McHenry, 2005). In addition, the ε subunit has an internal CBM, which is responsible for the replication-promoting role of the ε subunit (Jergic et al., 2013; Studwell and O’Donnell, 1990; Toste Rêgo et al., 2013). During processive replication, the α and ε subunits of the Pol III complex occupy both clefts of a β_2_ clamp molecule (Fernández-Leiro et al., 2015; Jergic et al., 2013; Toste Rêgo et al., 2013). However, unlike the internal CBM of the α subunit, the CBM of the ε subunit has a relative low binding affinity to the cleft; this results in its frequent detachment from a cleft, causing temporary pauses during processive replication (Loparo et al., 2011; J. Yang et al., 2004). Given the dynamic nature of the ε subunit-β_2_ clamp interaction, it may constitute a key factor in regulating the access of cleft-binding proteins, such as TLS polymerases.

Here we show that dynamic interactions between Pol III and the β_2_ clamp dictate the fate of lesion-stalled replisomes. We demonstrate that when Pol IV is present at optimal levels, TLS occurs at the fork through sequential polymerase switching between Pol III and Pol IV. In contrast, when Pol IV is present at suboptimal levels, a higher fraction of lesion-stalled replisomes are resolved by repriming replication downstream, leaving a ssDNA gap. We show that besides its canonical proofreading function, the ε subunit plays a gatekeeping role that largely prevents Pol IV from replacing Pol III during processive replication and limits the usage of TLS to resolve lesion-stalled replisomes. Central to this gatekeeping role is the dynamic interaction between the ε subunit and the β_2_ clamp. When this interaction is strengthened *in vitro*, Pol IV-mediated TLS at the fork is suppressed, leading to more lesion skipping. Conversely, when the ε-β_2_ interaction is weakened, Pol IV more efficiently mediates TLS at the fork and processive replication is more readily inhibited by Pol IV. In cells, when the SOS-response is not induced, TLS by all three TLS polymerases is minimally employed to resolve lesion-stalled replisomes. However, weakening the ε-β_2_ clamp interaction leads to resolution of a much larger fraction of lesion-stalled replisomes through TLS, indicating that the strength of the ε-β_2_ interaction is tuned to suppress TLS in uninduced cells. Collectively, these results show that the ε-β_2_ interaction sets a kinetic barrier to the access of TLS polymerases to the extending primer and thus suppresses TLS.

## Results

### Rapid and efficient Pol IV-mediated TLS within lesion-stalled replisomes

In an effort to determine how processive replication and TLS are coordinated, we reconstituted Pol IV-mediated TLS on a rolling circle DNA template that contains a site-specific N^2^-furfuryl dG (N^2^-FFdG) lesion on the leading strand template (Figures 1A and S1A). N^2^-FFdG is an attractive model lesion because Pol IV is proficient at replicating past N^2^-dG adducts and structurally related DNA lesions can be created in living cells by treatment with nitrofurazone (NFZ) (Jarosz et al., 2006). In rolling circle replication, replication proceeds over the circular template multiple times, generating a long leading strand tail that serves as a template for discontinuous lagging strand Okazaki fragments (Figure 1A) (Wu et al., 1992a; 1992b; Zechner et al., 1992a; 1992b). Replication of the lesion-free control template by the reconstituted *E. coli* replisome resulted in the rapid formation of replication products that can be resolved by denaturing gel electrophoresis: a resolution-limited leading strand product band and a distribution of smaller lagging strand products along with a fraction of unreplicated templates (Figures 1B and S1B) (Wu et al., 1992a). The resolution limit of our gel is approximately 45 kilonucleotides, and therefore accumulation of leading strand products at the resolution limit indicates that at least 6 cycles of rolling circle replication occurred on each template. Consistent with a prior observation that an N^2^-FFdG blocks primer extension by Pol III α (Kath et al., 2014), replication of the N^2^-FFdG-containing template in the absence of Pol IV was strongly attenuated by a single N^2^-FFdG on the leading strand template (Figure 1B). We also observed faint, discrete bands between resolution-limited replication products and unreplicated templates (Figures 1B and 1C). As these products were created only in the presence of N^2^-FFdG, they represent lesion-stalled replisomes that have undergone different multiples of replication around the template with TLS over the lesion inefficiently mediated by Pol III (Figures1B-D) (Nevin et al., 2017).

**Figure 1.**
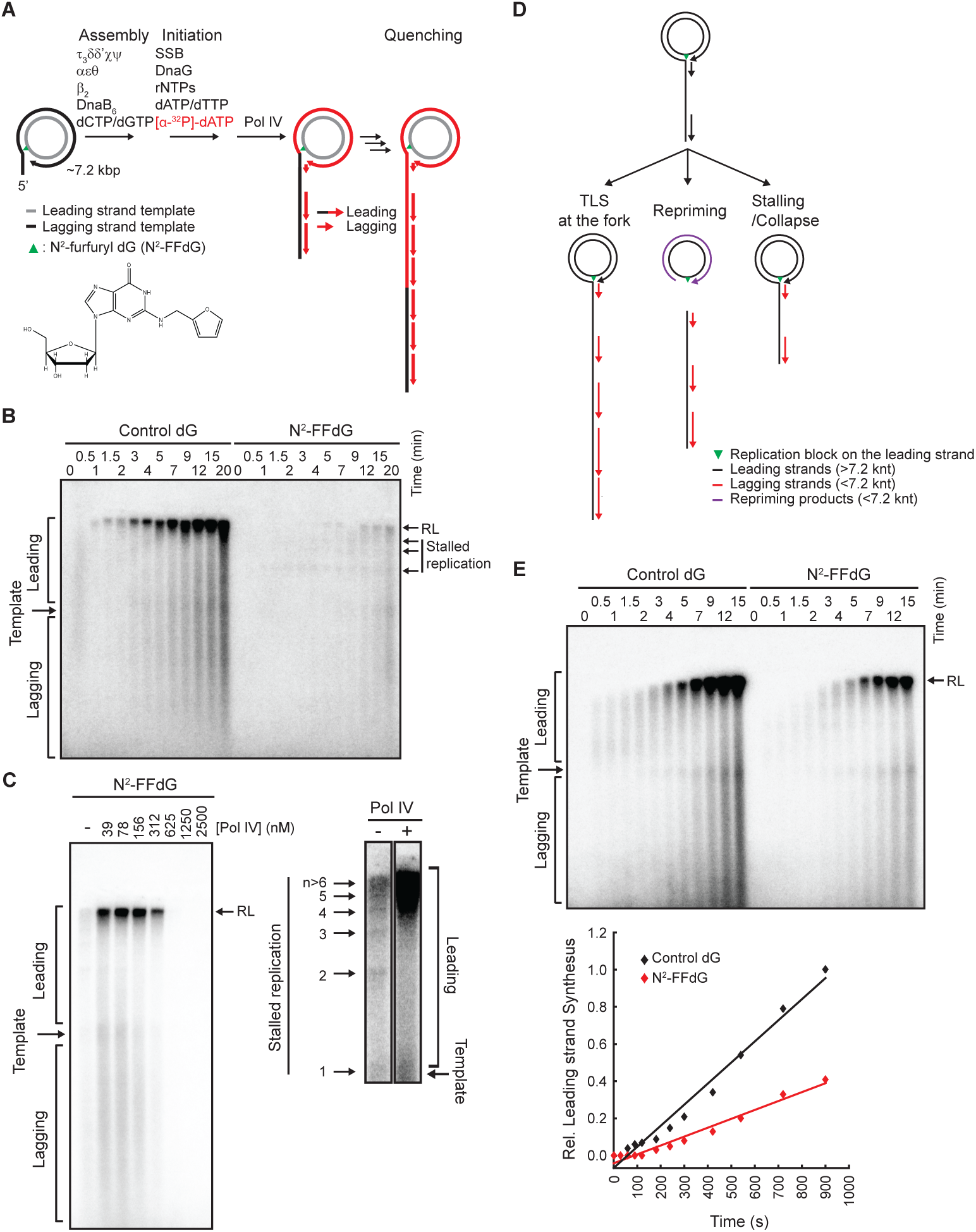
*In vitro* reconstitution of Pol IV-mediated TLS. A. A rolling-circle replication-based TLS assay. Replication reactions were performed through the indicated steps. Newly synthesized DNAs were labeled with incorporated [^32^P]-dATP during replication, separated on a denaturing gel and visualized by autoradiography. B. A single N^2^-FFdG in the leading strand template potently inhibits the formation of resolution-limited (RL) leading strand products by the *E. coli* replisome. Control, a lesion-free template; N^2^-FFdG, a N^2^-FFdG-containing template. C. Pol IV promotes replication of the N^2^-FFdG-containing template. (Left) Replication of N^2^-FFdG-containing template in the presence of indicated amounts of Pol IV. Reactions were quenched 12 minutes after initiation. (Right) Magnified view of the same gel. -, no Pol IV; +, 156 nM Pol IV; numbers (n), number of passages through a N^2^-FFdG lesion. D. A cartoon depicting the fates of lesion stalled replisomes in the rolling-circle assay and their associated replication products. E. Pol IV-mediated TLS at the fork only modestly slows replication of the N^2^-FFdG-containing template. 100 nM Pol IV was included in replication of both the control and N^2^-FFdG-containing templates. The lines are linear regressions of integrated intensities of leading strand products (see “Methods and Materials” section).

To determine whether Pol IV-mediated TLS might resolve this stalling, we examined the effect of Pol IV on replication of the N^2^-FFdG-containing template. Upon addition of increasing amounts of Pol IV, synthesis of both the leading and lagging strands was gradually restored and long leading strand replication products accumulated at the resolution limit before replication was fully inhibited by Pol IV at high concentrations (Figures 1C and S1E) (Indiani et al., 2009). The robust formation of these resolution-limited products shows that TLS occurs efficiently over the lesion at the fork as repriming of leading strand synthesis would result in a gap that terminates rolling circle replication. This termination results from the displacement of the circular leading strand template that occurs when the helicase runs into the strand discontinuity resulting from repriming (Figure 1D).

Consistent with the requirements of Pol IV-mediated TLS in cells, the ability of Pol IV to promote replication of the N^2^-FFdG-containing template required both its catalytic and clamp-binding activities (Figures S1C-F) (Becherel et al., 2002). Pol IV-mediated TLS occurred less efficiently over an N^2^-(1-carboxyethyl)-2’-deoxyguanosine (N^2^-CEdG)-lesion and was strongly blocked by tetrahydrofuran (THF), indicating that our rolling-circle assay is sensitive to the efficiency by which Pol IV mediates TLS over various lesions (Figure S1G) (Maor-Shoshani et al., 2003; B. Yuan et al., 2008).

Moreover, at optimal Pol IV concentrations, the rate of replication of the N^2^-FFdG-containing template, as measured by quantifying leading strand replication products (Figure S1C), was only slowed by ∼50% compared to replication of the control template (Figure 1E). Given that the polymerization rate of the Pol IV-based *E. coli* replisome is ∼10 nt/s (Indiani et al., 2009; Kath et al., 2014), which is over 50 times lower than that of the Pol III-based replisome, these results indicate that Pol IV only briefly switches with Pol III (Kath et al., 2014; Wagner et al., 2000). Collectively these results demonstrate that within our reconstitution, Pol IV can efficiently mediate TLS at the replication fork by switching with Pol III, synthesizing a small patch of DNA over the lesion.

### The ε-cleft contact mediates polymerase exchange during TLS

As Pol III core (αεθ) occupies both clefts of the β_2_ clamp during processive replication (Jergic et al., 2013) and Pol IV must bind to a cleft for its action (Figures S1E and S1F), we sought to determine how Pol III disengages from the clamp to allow for Pol IV to mediate TLS at the fork. To address this question, we varied the strength of the interaction between Pol III core and the β_2_ clamp and examined the effect on TLS. The α and ε subunits of Pol III core (αεθ) each occupy a cleft via independent clamp-binding motifs (CBMs), albeit with vastly different affinities; the ε-cleft contact is ∼250 fold weaker than the α-cleft interaction (Figure 2A) (Dohrmann and McHenry, 2005; Jergic et al., 2013). Replacing the wild-type CBM of the ε subunit with a mutant CBM (ε_L_), which binds the cleft ∼500 times tighter than the wild-type CBM (Jergic et al., 2013), suppressed Pol IV-mediated TLS compared with the wild-type Pol III core (αεθ) and required higher Pol IV concentrations for optimal TLS (Figure 2B). Normalizing replication of the lesion containing template by replication of the lesion-free control template (see Methods) showed that the *ε*_*L*_ mutation suppressed Pol IV-mediated TLS to ∼40% (Figure 2B inset, right panel). Strengthening the interaction also modestly enhanced replicative activity on a lesion-free template (Figure 2B inset, left panel) (Jergic et al., 2013). These results suggest that the disengagement of the ε subunit from the β_2_ cleft at a lesion is necessary for Pol IV to bind to the β_2_ clamp and perform TLS.

**Figure 2.**
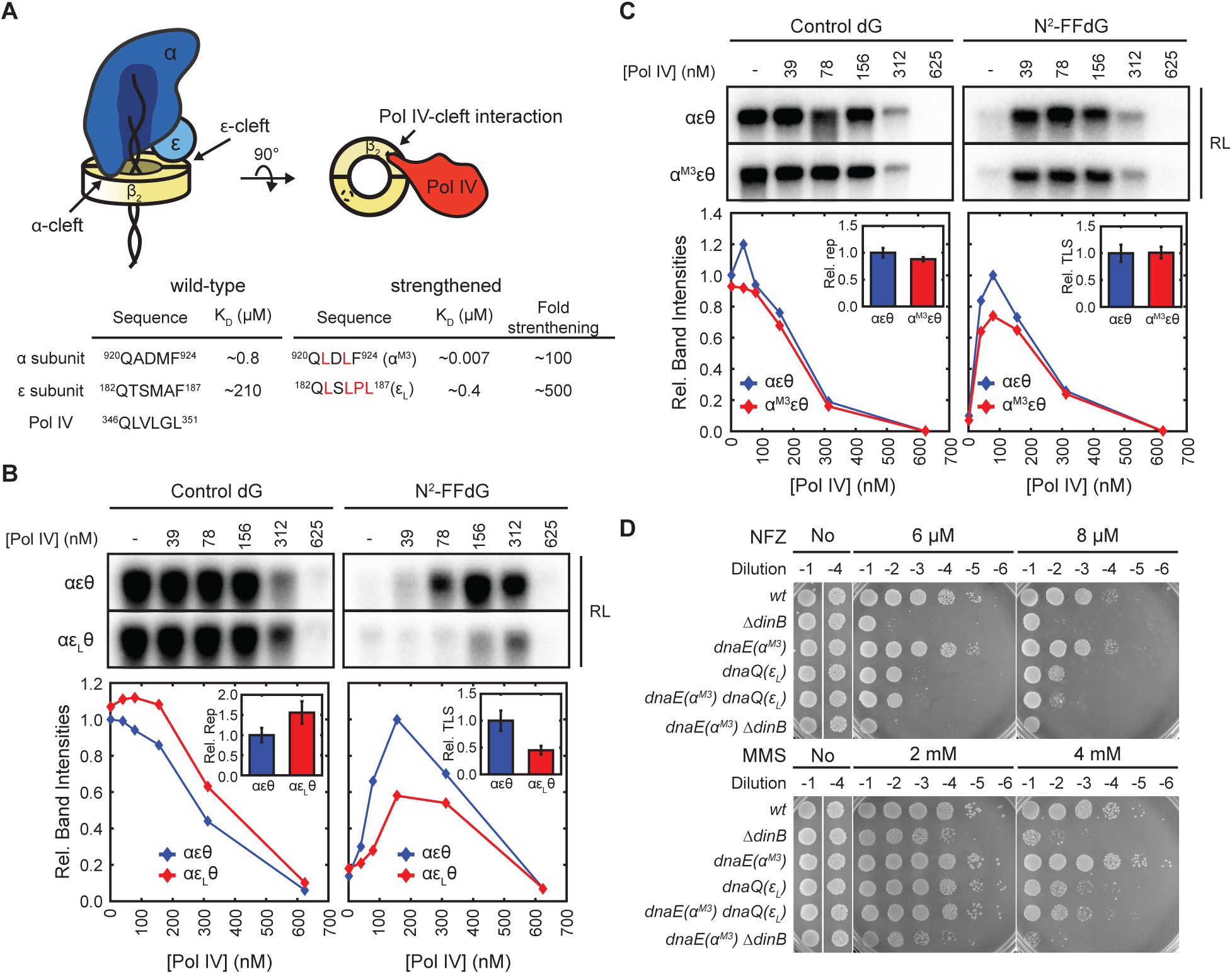
ε-cleft mediates polymerase exchange. A. Interactions between Pol III core (αεθ) or Pol IV and the β^2^ clamp via the clamp binding motif (CBM)–cleft interaction. (Top) α- cleft, the cleft occupied by the α subunit through its internal CBM. ε-cleft, the cleft occupied by the ε subunit through its internal CBM. Pol IV interacts with a cleft via the C-terminal CBM. For simplicity, the θ subunit of Pol III core is not depicted. (Bottom) Mutations on CBMs to strengthen the interaction with a cleft. B. Strengthening the ε-cleft interaction suppresses *in vitro* TLS. (Top) Resolution-limited (RL) leading strand replication products. (Bottom) Relative band intensities were calculated with respect to the band intensity in the absence of Pol IV for the lesion-free template and the maximal band intensity for the lesion containing templates in the presence of Pol IV, respectively. αεθ, wild-type ε- containing replisome; αε_L_θ, ε_L_-containing replisome. (Inset, left) Relative replication (Rel. Rep) corresponds to replication of the control template normalized by replication of the αεθ-containing replisome at [Pol IV] = 156 nM. (Inset, right) Relative TLS (Rel. TLS) was calculated by first normalizing replication of the lesion-containing template with replication of the control and then calculating the ratio with respect to the αεθ replisome. Reported values correspond to [Pol IV] = 156 nM (mean ± SD, n = 3) C. Enhancing the α-cleft contact has little effect on *in vitro* replication or TLS. (Top) Resolution-limited (RL) leading strand replication products. (Bottom) Integrated intensities of leading strand replication products. αεθ, wild-type ε-containing replisome; α^M3^εθ, α^M3^-containing replisome. (Inset, left) Relative replication (Rel. Rep), replication of the control template at [Pol IV] = 156 nM. (Inset, right) replication-normalized relative TLS (Rel. TLS) at [Pol IV] = 156 nM (mean ± SD, n > 2) D. Strengthening the ε-cleft interaction sensitizes cells to NFZ and MMS. Cultures of indicated strains were serially diluted and spotted on LB-agar plates containing indicated concentrations of either NFZ or MMS. *dnaQ* and *dnaE*, the genes encoding the and the subunits respectively.

As the α subunit also contains a CBM, we next considered if disengagement of α is additionally required for Pol IV-mediated TLS (Figure 2A). To address this possibility, we examined the effect of strengthening the α-cleft interaction within our biochemical reconstitution. Unlike the effects of strengthening the ε-cleft interaction, replacing the wild-type CBM of the α subunit with a mutant CBM (α^M3^), which binds the cleft ∼100 times tighter than the wild-type CBM (Dohrmann and McHenry, 2005) – even ∼50 times tighter than the ε_L_-cleft interaction - had little effect on replication of the lesion-free or lesion-containing template (Figure 2C). Collectively, these results demonstrate that disengagement of only the ε subunit, not the α subunit, within lesion-stalled replisomes promotes Pol IV to bind to a cleft and perform TLS.

To validate these *in vitro* observations in cells, we introduced the *dnaQ(ε*_*L*_) and *dnaE(α*_*M3*_) mutations, individually or in combination, into their respective genomic loci and examined their effects on cellular sensitivity to the DNA damaging agents NFZ and MMS. Cells lacking Pol IV (*ΔdinB*) were strongly sensitized to NFZ and methyl methanesulfonate (MMS), indicating that Pol IV-mediated TLS contributes to cellular tolerance of these agents (Figure 2D). Consistent with our *in vitro* observations, the *dnaQ(ε*_*L*_) strain was sensitized to both NFZ and MMS whereas the *dnaE(α*^*M3*^) strain retained wild-type tolerance (Figure 2D). Furthermore, a strain containing both the *dnaQ(ε*_*L*_) and *dnaE(α*^*M3*^) mutations (*dnaQ(ε*_*L*_*) dnaE(α*^*M3*^)) resembled the sensitivity of *dnaQ(ε*_*L*_), indicating that the strengthened ε-cleft interaction is responsible for the increased sensitivity. This increased sensitivity of the *dnaQ(ε*_*L*_) strain is due to defective Pol IV-mediated TLS because *dnaQ(ε*_*L*_) was epistatic to *ΔdinB* (Figure S2A). The *dnaQ(ε*_*L*_) and *dnaE(α*^*M3*^) strains grew normally and retained nearly wildtype DNA content (Figures S2B and S2C). Furthermore, the *dnaQ(ε*_*L*_) strain retained similar numbers of replisome foci to the *dnaQ*^*+*^ strain both in untreated and NFZ-treated cells (Figure S2D and S2E). The *dnaQ(ε*_*L*_) and *dnaE(α*^*M3*^) strains also retained nearly wild-type SOS-responses to both NFZ and MMS. Collectively these results rule out the possibility that the increased sensitivity of the *dnaQ(ε*_*L*_) strain resulted from a general defect in DNA replication or the DNA damage response.

### ε subunit acts as a gatekeeper

As the *dnaQ(ε*_*L*_) mutation stimulates proofreading of Pol III core (αεθ) (Park et al., 2017; Toste Rêgo et al., 2013), we next addressed whether the ε_L_–containing replisome suppresses TLS through futile cycles of TLS and proofreading, impeding the transition from TLS to processive replication (Figure 3A) (Fujii and Fuchs, 2004; Jarosz et al., 2009). If the proofreading function of the ε subunit counteracts Pol IV-mediated TLS, abrogating the exonuclease activity of the ε subunit should promote Pol IV-mediated TLS. Indeed, when the catalytic residues of the ε subunit were mutated (αε^D12A,E14A^θ) (Fijalkowska and Schaaper, 1996), Pol IV-mediated TLS was promoted (Figure 3B), indicating that the proofreading function of the ε subunit antagonizes Pol IV-mediated TLS over N^2^-FFdG, likely through futile cycles. In contrast to the effect on TLS, abrogating the exonuclease activity of the ε subunit reduced replication on the control template (Figure 3B), indicating that exonuclease activity facilitates processive replication. Importantly, we found that the *dnaQ(ε*_*L*_) mutation still strongly suppressed Pol IV-mediated TLS within the exonuclease-dead *dnaQ*-containing replisome (αε^D12A,E14A^θ), which was comparable to the suppression within the wild-type replisome (αεθ) (Figure 3B). This result indicates that strengthening the ε-cleft interaction suppresses Pol IV-mediated TLS by inhibiting the interaction of Pol IV with the β_2_ clamp rather than promoting futile cycles of TLS and proofreading.

**Figure 3.**
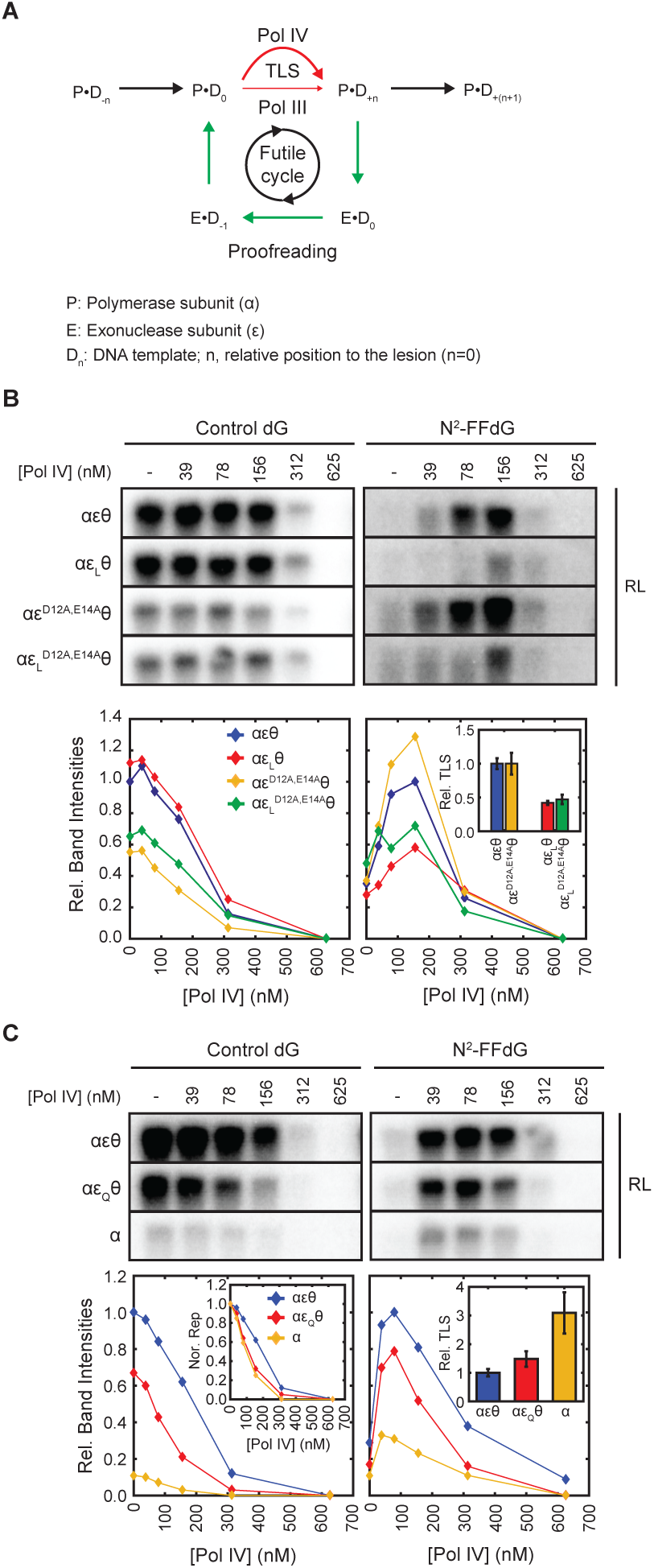
ε subunit suppresses TLS through a gatekeeping role. A. Proofreading can lead to abortive TLS through futile cycles. Processive replication (black thick arrow); Pol IV-mediated TLS (red thick arrow, fast); Pol III-mediated TLS (red thin arrow, slow); Proofreading (green arrows) B. Strengthening the ε-cleft interaction with the *dnaQ(ε*_*L*_) mutation suppresses TLS at the fork for both wild-type and catalytically defective ε. (Top) Resolution-limited (RL) leading strand replication products. (Bottom) Integrated intensities of these leading strand replication products. ε^D12A,E14A^, catalytically defective ε; ε^LD12A,E14A^, catalytically defective ε with the *dnaQ(ε*_*L*_) mutation. (Inset, right) Replication-normalized relative TLS at [Pol IV] = 78 nM (mean ± SD, n = 2). C. Weakening the ε-cleft interaction promotes Pol IV-mediated TLS at the fork. Resolution-limited (RL) leading strand replication products resulting from replication of a lesion-free control template (Top, left) or a N^2^-FFdG-containing template (Top, right) by the indicated Pol III complexes. (Bottom) Integrated intensities of these leading strand replication products. (Inset, left) Inhibition of replication by Pol IV; replication is normalized to replication in the absence of Pol IV. (Inset, right) Replication-normalized relative TLS at [Pol IV] = 156 nM (mean ± SD, n > 2). αεθ, wild-type replisome; αε^Q^θ, ε^Q^-containing replisome; α, ε-free replisome.

Given these results suggest an exonuclease activity-independent gatekeeping role of the ε subunit, we hypothesized that weakening the ε-cleft interaction or removing ε altogether would make replication more potently inhibited by Pol IV and increase Pol IV-mediated TLS. To weaken the ε-cleft interaction, we introduced the *dnaQ(ε*_*Q*_) mutation, which substantially decreases affinity for the clamp (K_D_ >2 mM), (Jergic et al., 2013). Indeed, the ε_Q_-containing replisome (αε_Q_θ) exhibited reduced replicative activity on the control template and replication was more potently inhibited by Pol IV as compared with the wild-type replisome (αεθ) (Figure 3C, left inset). On the N^2^-FFdG-containing template, Pol IV-mediated TLS occurred more efficiently in the ε_Q_-containing replisome (αε_Q_θ) compared with the wild-type replisome (αεθ) when normalized for replication on the control template (Figure 3C, right inset). Removal of the ε subunit further increases the efficiency and potency of Pol IV mediated TLS, suggesting that in the absence of the ε subunit (Figure 3C, α alone replisome), Pol IV may frequently bind to the β_2_ clamp even during processive replication. Similar observations were also made with the N^2^-CEdG-containing template (Figure S3A). These results demonstrate that it is a gatekeeping activity of the ε subunit rather than its exonuclease activity that limits the access of Pol IV and potentially other clamp binding proteins to lesion-stalled replisomes.

### Strength of the ε-cleft interaction determines pathway choice between TLS at the fork and repriming

Given prior observations that the reconstituted *E. coli* replisome can reprime leading strand synthesis (Yeeles and Marians, 2013; 2011), we next asked if suppression of TLS at the fork by strengthening the ε- cleft interaction increases repriming. To determine whether the N^2^-FFdG-stalled replisome reprimed downstream of the lesion, we used a Southern blot to detect the expected repriming products (Figures 1D, and 4A and 4B). In the absence of Pol IV, replication of the N^2^-FFdG-containing template created replication products that were detected with leading strand specific Southern blot probes (Figure 4A and 4B, and S4A and S4B) (Wu et al., 1992a). Replication products that ran as discrete bands above the template were leading strand replication products that resulted from processive rolling circle replication because these were detected only with leading strand probes and were longer than the template (No Pol IV in Figures 4B, and S4A and S4B). Additionally, we detected diffuse replication products that ran below the template. These replication products were not created in the absence of DnaG, but unlike Okazaki fragments these were detected with leading strand probes (No Pol IV in Figures 4B, and S4C and S4D). These short leading strand products were better detected with a distal leading strand probe (1901 nt probe) than a proximal leading strand probe (40 nt probe) (Figures 4A and S4E), consistent with prior reports that repriming occurs a few hundred nucleotides downstream of the lesion (Yeeles and Marians, 2013). Collectively, these results indicate that replisomes stalled at N^2^-FFdG reprime DNA synthesis downstream of N^2^-FFdG, leaving a single-stranded DNA gap between the lesion and a newly synthesized leading strand RNA primer (Figure 4A).

**Figure 4.**
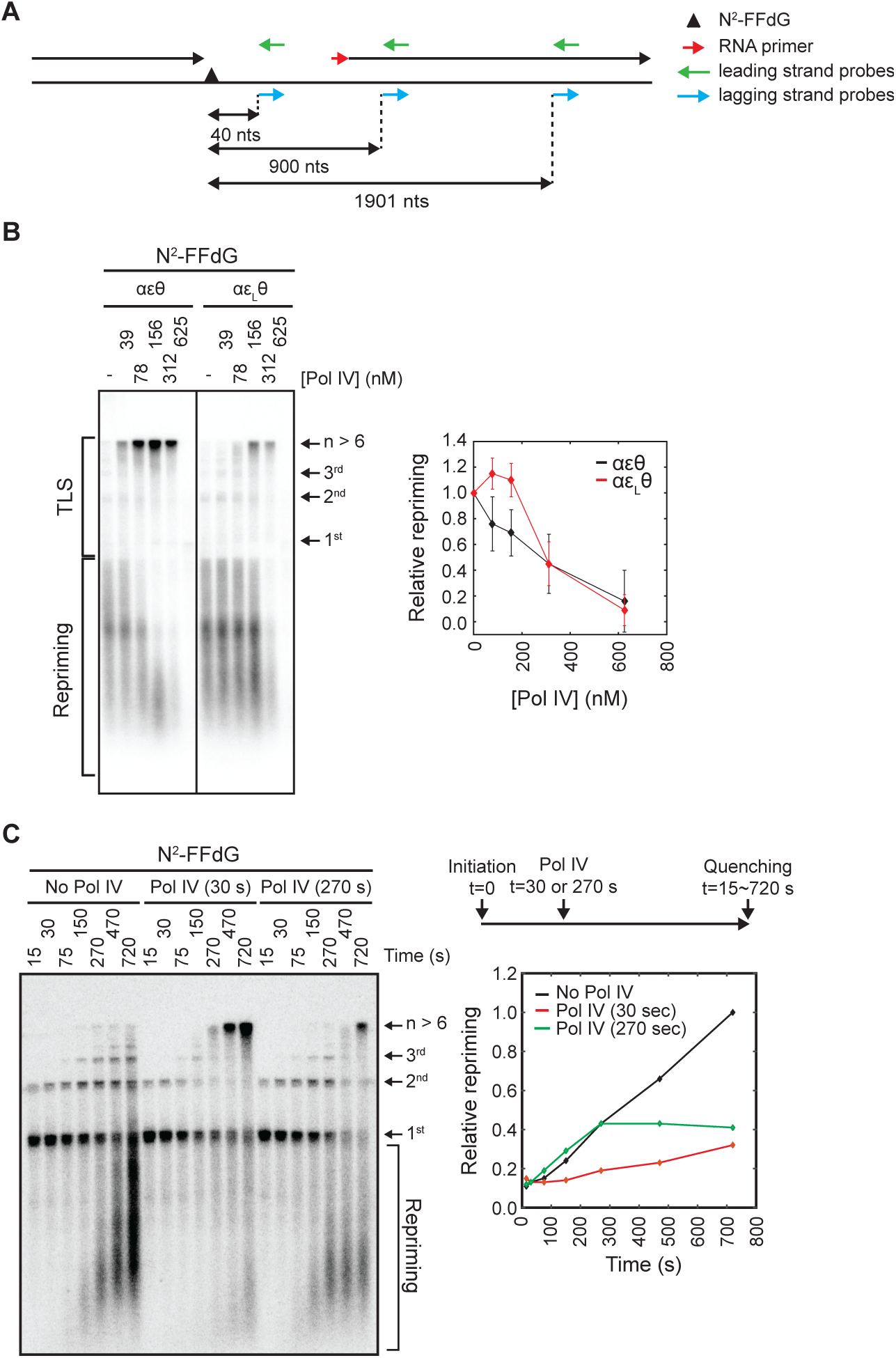
Pol IV diverts lesion-stalled replisomes from repriming to TLS. A. Southern blot probes used in this study. A lesion-stalled replisome on the leading strand template reprimes downstream of the lesion, leaving a ssDNA gap between the lesion and a newly synthesized downstream primer. B Strengthening of the -cleft interaction suppresses TLS at the fork resulting in persistent repriming. (Left) Lesion-stalled replisomes reprime downstream of N^2^-FFdG. Repriming and leading strand replication products resulting from replication of the N^2^-FFdG-containing template by wild-type (αεθ) or ε_L_-containing (αε_L_θ) replisomes. Replication products were separated in a denaturing agarose gel and detected by Southern blot with a mixture of leading strand probes (900 and 1901 nts) shown in Figure 4A. (Right) Integrated intensities of repriming replication products (mean ± SD, n = 3). C. Pol IV-mediated TLS at the fork kinetically competes with repriming. (Right, top) Replication of the N^2^-FFdG-containing template was initiated in the absence of Pol IV, and Pol IV (100 nM final) was added to the reaction 30 or 270 seconds after initiation. (Left) Reactions were quenched at the indicated time after initiation, and replication products were separated in a denaturing agarose gel and detected by Southern blot with a leading strand probe (900 nts) shown in Figure 4A. (Right, bottom) Quantitation of repriming products. Amounts of repriming products were plotted relative to the amount of repriming products detected at the final time point (720 seconds) in the absence of Pol IV.

The addition of increasing concentrations of Pol IV to replication reactions of the N^2^-FFdG-containng template led to a gradual increase in TLS at the fork and a concomitant decrease in repriming, suggesting that Pol IV-mediated TLS competes with repriming (Figures 4B and S4F-H). Indeed, when wild-type Pol IV (100 nM) was added with varying time delays after the initiation of replication, stalled replisomes were rapidly released into the TLS pathway and no further increase in repriming was observed (Figure 4C).

Notably, for the ε_L_-containing replisome, while TLS was suppressed, repriming persisted in the presence of higher concentrations of Pol IV as compared with the wild-type replisome (Figure 4B). To examine whether strengthening the ε-cleft interaction increases the intrinsic propensity of the replisome to reprime, we compared the time course of repriming in the absence of Pol IV between the wild-type and ε_L_-containing replisomes. Both replisomes had a similar partitioning between Pol III-mediated TLS over the lesion and repriming with repriming occurring at similar rates (t_1/2,apparent_ ∼ 4 minutes) (Figure S4I). Therefore, the persistent repriming of the ε_L_-containing replisome at higher concentrations of Pol IV most likely results from suppression of Pol IV-mediated TLS.

### *In vivo*, Pol IV-mediated TLS increases upon weakening the ε-β_2_ clamp interaction

Our *in vitro* observations demonstrate that the ε-cleft interaction plays a decisive role in determining the fate of lesion-stalled replisomes. To test whether this also happens in living cells, we employed an *in vivo* assay that quantitatively measures the fraction of stalled replisomes that are resolved either by TLS or via damage avoidance (DA) pathways at a site-specific lesion located in the chromosome. The experimental system (Figures 5A and S5A), which is based on phage lambda site-specific recombination, consists of two major components: a recipient *E. coli* strain with a single *attR* site and a non-replicating plasmid construct containing the single lesion of interest, an *attL* site and an ampicillin resistance gene. A site-specific recombination reaction between *attL* and *attR* is controlled by ectopic expression of phage lambda integrase and excisionase proteins (Valens et al., 2004) and leads to the integration of the lesion-containing plasmid into the chromosome. Integrants are selected on the basis of their resistance to ampicillin. The chromosomal region where integration takes place carries the 3’-end of the *lacZ* gene fused to *attR*, (min 17’ in the *E. coli* chromosome) while the remaining 5’-end is located on the incoming plasmid fragment in fusion with *attL*. Precise integration restores a functional β-galactosidase gene (*lacZ*^*+*^). A short sequence heterology inactivates the *lacZ* gene in the “opposite” strand across from the single lesion and serves as a genetic marker to allow strand discrimination during replication (Veaute and Fuchs, 1993). The recipient strain carries both *uvrA* and *mutS* deletions to prevent the single lesion and the sequence heterology from being repaired by nucleotide excision repair and mismatch repair, respectively. Functional LacZ is expressed only when the lesion-containing leading strand template is bypassed by TLS events with the exception of events that contain a frameshift mutation. Therefore, colonies resulting from TLS events appear as blue-sectored colonies on X-Gal containing plates (Figures 5A and S5A) (Pagès et al., 2012). On the other hand, white colonies result from recA-mediated Homology-Dependent Gap Repair (Laureti et al., 2015)

**Figure 5.**
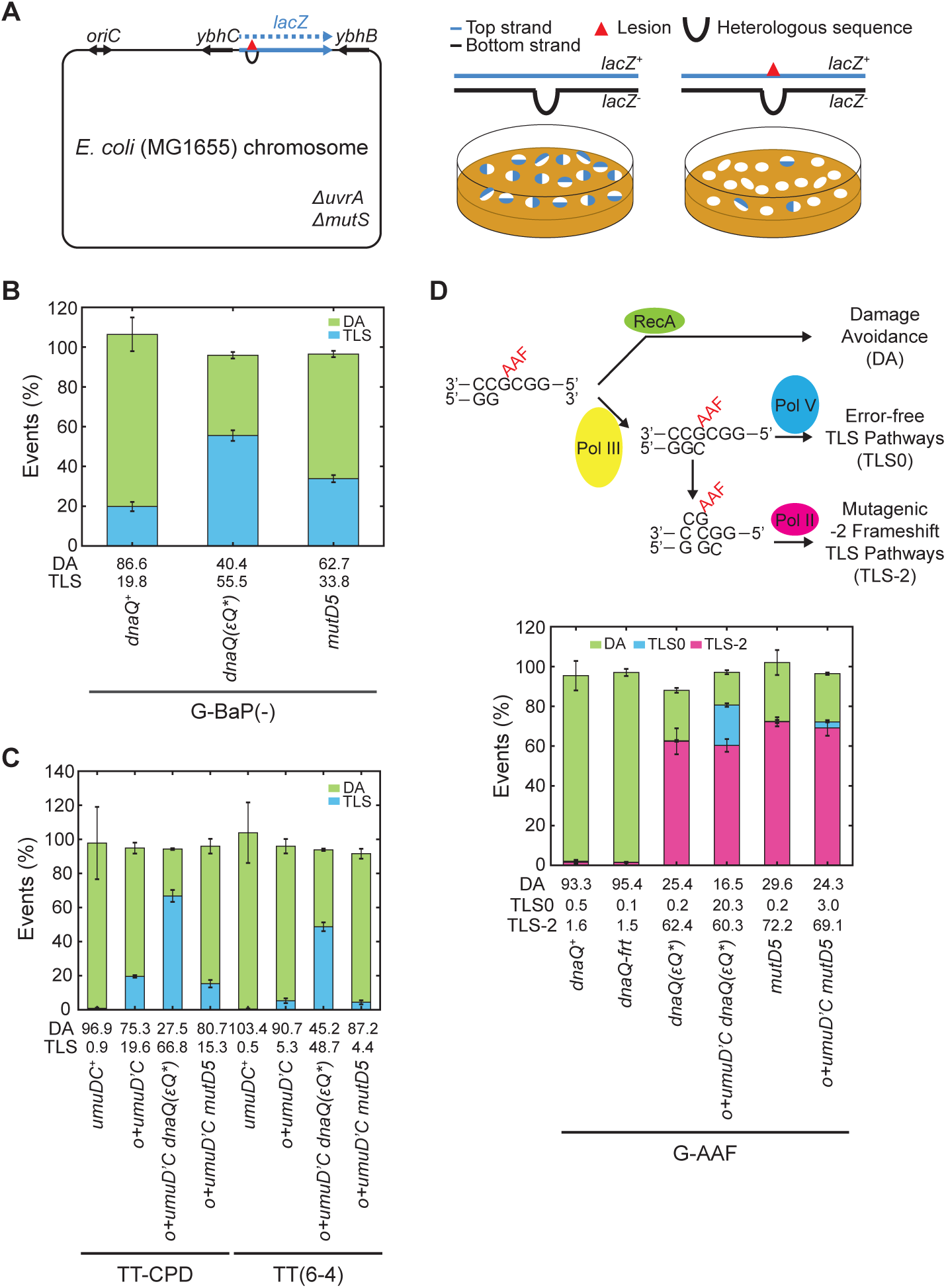
Weakening the ε-cleft interaction increases utilization of TLS in cells. A. A schematic diagram of the *in vivo* TLS assay. Following integration of lesion-free lacZ^-^/lacZ^+^ heteroduplex constructs into the genome, only blue-sectored colonies are formed. In contrast, when a single replication-blocking lesion is integrated, only a fraction of colonies formed are blue-sectored. The sectored colonies represent TLS events as explained in detail in the main text. Recipient strains in B-D with various *dnaQ* and *umuDC* alleles are described in the main text. TLS0 and TLS-2 are TLS events without frameshift and −2 frame shift respectively; DA, damage avoidance. B. Extent of Pol IV-mediated TLS across G-BaP(-) (N^2^-(-)-trans-anti-benzo[a]pyrene dG) in indicated strains. C. Extent of Pol V-mediated TLS across TT-CPD and TT(6-4) in indicated strains. D. Pol II- and Pol V-mediated TLS across G-AAF (N^2^-acetylaminofluorene dG) in indicated strains. (Top) Scheme of Pol II- or Pol V-mediated TLS across G-AAF. (Bottom) Extent of Pol II- and Pol V-mediated TLS over G-AAF in indicated strains. Each data point represents the average of at least four independent experiments and error bars indicate the standard deviations.

To investigate Pol IV-mediated TLS, we introduced a single (−)-trans-anti-benzopyrene-N^2^-dG adduct (dG-BaP(−)) into the *E. coli* genome, known to be bypassed by Pol IV alone (Seo et al., 2006; Shen et al., 2002). In the *dnaQ*^*+*^ background, the majority of G-BaP(-)-stalled replisomes were processed via DA pathways, while only 20% were resolved by TLS (Figure 5B). However, in the *dnaQ(ε*_*Q*_*\*)* background, which weakens the ε-cleft interaction, 55% of the stalled replisomes were resolved by TLS with a corresponding drop in the use of DA pathways. Since the basal SOS response in the *dnaQ(ε*_*Q*_) strain was comparable to that in the *dnaQ*^*+*^ strain (Figure S5B), we ruled out the possibility that the increase in Pol IV-mediated TLS in the *dnaQ(ε*_*Q*_*\*)* strain resulted from elevation of Pol IV expression levels.

As the *dnaQ(ε*_*Q*_*\*)* mutation also reduces the processivity of proofreading (Park et al., 2017; Toste Rêgo et al., 2013), we examined if the increase in TLS in *dnaQ(ε*_*Q*_*\*)* background was primarily due to a decrease in proofreading activity. To test this possibility, we abolished the exonuclease activity of the ε subunit with the *mutD5* mutation (Supplementary table 1) (Degnen and Cox, 1974; Maruyama et al., 1983) and assessed the fraction of DA and TLS events over the G-BP(-) lesion. In the *mutD5* background, participation of TLS increased from ∼20 to ∼34 % but still remained significantly below the 55% value seen in the *dnaQ(ε* _*Q*_^*\**^) strain (Figure 5B). Therefore, the additional increase in TLS seen upon weakening the ε-cleft contact supports the model that the ε subunit plays a gatekeeping role in regulating access of Pol IV during TLS in living cells.

### The ε subunit acts as a gatekeeper for all three *E. coli* TLS polymerases

Next, we asked whether the ε-cleft interaction plays a similar gatekeeping role in Pol V-mediated TLS across TT-CPD and TT(6-4) lesions. In the *dnaQ*^*+*^ background, replisomes stalled at TT-CPD and TT(6-4) were fully processed by the DA pathway as no Pol V-mediated TLS was measured. This lack of TLS is likely due to the absence of the active form of Pol V (UmuD’_2_C) as UmuD is not processed into UmuD’ under non-SOS induced conditions (Figure 5C) (Fuchs and Fujii, 2013). In order to be able to observe Pol V-mediated TLS, we engineered a strain that carries *umuD’* instead of *umuD* at the original *umuDC* locus (*umuD’C* strain); this strain constitutively expresses the active form of Pol V (Ennis et al., 1995). In this background, Pol V-mediated TLS corresponded to ∼20% and ∼5 % for TT-CPD and TT(6-4) lesions, respectively (Figure 5C). Introduction of the *dnaQ(ε*_*Q*_*\*)* allele in the *umuD’C* background further stimulated TLS to ∼70 and ∼50 % for TT-CPD and TT(6-4), respectively (Figure 5C). In contrast, introducing the proofreading-deficient *mutD5* allele in the *umuD’C* background, had no effect on the level of TLS, suggesting that the robust increase in TLS in the *dnaQ(ε*_*Q*_*\*)* strain resulted from an increased accessibility of Pol V to the clamp rather than from a defect in proofreading (Figure 5C).

Finally, to investigate Pol II-mediated TLS, we introduced a N^2^-acetylaminofluorene dG (G-AAF) lesion in the *NarI* sequence context; this lesion can be bypassed by either Pol II- or Pol V-mediated TLS leading to −2 or error-free bypass, respectively (Figure 5D) (Fuchs and Fujii, 2007; Napolitano et al., 2000). In contrast to what was observed for Pol IV and Pol V cognate lesions, Pol II-mediated TLS over the G-AAF lesion was comparably stimulated in the *dnaQ(ε*_*Q*_*\*)* and *mutD5* strains (Figure 5D). The similar levels of TLS in the *dnaQ(ε*_*Q*_*\*)* and *mutD5* backgrounds are most likely due to the peculiar mechanism of Pol II-mediated −2 frameshift mutagenesis, which involves elongation of a “slipped lesion terminus” (Figure 5D) (Fuchs and Fujii, 2007; Fuchs and Napolitano, 1998). As revealed in the crystal structure, Pol II is uniquely suited to elongate this frameshift intermediate (Wang and W. Yang, 2009). Upon stalling at the G-AAF lesion, Pol III dissociates from the P/T junction, allowing for the formation of a −2 slippage intermediate; this intermediate is thermodynamically-favored by the stacking energy provided by insertion of the aromatic fluorene residue between the neighboring base pairs (Fuchs and Daune, 1972; Milhé et al., 1996). We speculate that in the *mutD5* strain the formation of the −2 slippage intermediate is enhanced due to a lack of Pol III-dependent proofreading, while in the *dnaQ(ε*_*Q*_*\*)* strain, Pol II has increased access to the clamp in order to extend this slippage intermediate.

With respect to error-free bypass of G-AAF by Pol V, TLS is stimulated (∼7 fold) in the *dnaQ(ε*_*Q*_*) strain compared to the *mutD5* strain (Figure 5D), in good agreement with the data observed for Pol V-mediated TLS across TT-CPD and TT(6-4) lesions. Productive TLS, requires Pol V to remain bound to the primer P/T junction in order to synthesize a TLS patch rather than to only insert a single nucleotide directly across from the lesion site (Fujii and Fuchs, 2004; Isogawa et al., 2018). As weakening the ε-cleft interaction in cells significantly increased the utilization of TLS over cognate lesions for all three TLS polymerases, we conclude that the ε-cleft contact acts as a gatekeeper to regulate access to the P/T junction.

## Discussion

### Conformational transitions of the replicative polymerase underlie pathway choice

During replication, replisomes encounter a diverse set of challenges that can stall replicative DNA polymerases, including DNA lesions and protein-DNA complexes (Marians, 2018; Merrikh et al., 2012). It remains poorly understood how replisomes select among distinct rescue mechanisms to cope with these different situations. In this study, we demonstrated that for replisomes stalled at leading strand lesions, the dynamic interaction between the exonuclease subunit (ε) of Pol III and the β_2_ clamp plays a crucial role in pathway choice between TLS at the fork and repriming (Figure 6A). We showed that the ε-cleft interaction acts as a steric gate to limit access of TLS polymerases to the primer/template (P/T) junction in a manner that is largely independent of its exonuclease activity.

**Figure 6.**
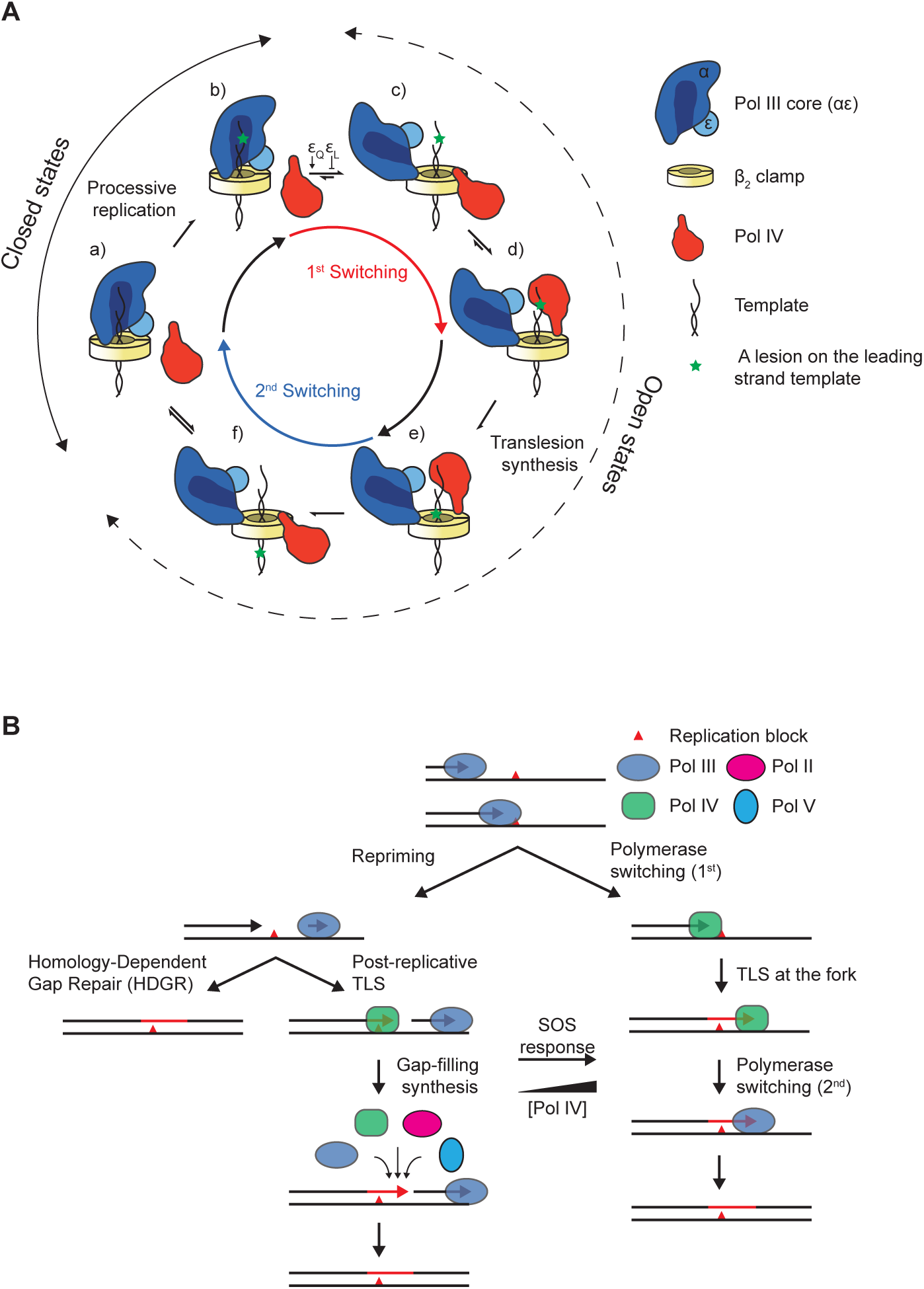
Conformational basis of pathway choice of lesion-stalled replisomes. A. Conformational transitions of Pol III core complex during TLS. a) Pol III core (αεθ) occupies both α- and ε-clefts during processive replication. b), c) Upon encountering a lesion on the leading strand, Pol III core swings away from the P/T junction while remaining bound to the β^2^ clamp through the α-cleft interaction. This conformational transition is either facilitated or suppressed by the *dnaQ(ε*_*Q*_) or the *dnaQ(ε*_*L*_) mutations, respectively. d),e) Pol IV binds to the ε-cleft and take over the P/T junction and performs TLS past the lesion. f) After synthesizing a short patch, Pol IV swings away from the P/T junction and Pol III core re-establishes the ε-cleft interaction and resumes processive replication. Within closed states, states a and b, the -cleft is occupied by the subunit. Within open states, states c though f, the -cleft is not occupied by the subunit. The θ subunit of Pol III core is not depicted for simplicity. B. Pathway choice for lesion-stalled replisomes during the SOS response: Repriming followed by gap-filling TLS vs TLS at the fork.

Pol IV gains access to the P/T junction by first binding to the cleft on the β_2_ clamp that is occupied by the ε subunit during processive replication (b through d in Figure 6A). Weakening the ε-cleft interaction (“opening the gate”, transition from b to c in Figure 6A) markedly increased TLS at the fork while only modestly decreasing processive replication. Strengthening the ε-cleft interaction (“closing the gate”, transition from c to b in Figure 6A) exerted the opposite effects. Therefore, the strength of this interaction is finely tuned to maximize processive replication on undamaged DNA while still enabling replisome remodeling by TLS polymerases upon the encounter of DNA damage.

Within stalled replisomes, recurring binding and unbinding between the ε subunit and the cleft form a kinetic barrier to Pol IV-clamp binding. Unbinding may be stimulated by a bulky lesion in the template strand; the inability of the narrow catalytic cleft of the replicative polymerase to accommodate the lesion-containing template could promote opening of the Pol III–clamp complex. Alternatively, replicating Pol III core complexes may sample the open state (reversible transition between b and c in Figure 6A) regardless of the presence of blocking lesions (Zhao et al., 2017). In either case, strengthening the ε-cleft interaction suppresses these conformational transitions and therefore TLS, and weakening the interaction has the opposite effect.

Recent observations demonstrate that the *E. coli* replisome readily exchanges Pol III holoenzyme from cytosol during processive replication (Beattie et al., 2017; Lewis et al., 2017; Q. Yuan et al., 2016), suggesting that Pol III core dissociation from the clamp may be required for Pol IV binding (Zhao et al., 2017). In contrast, our results suggest that Pol III core remains bound to the β_2_ clamp during Pol IV-mediated TLS as strengthening the α-cleft interaction had little effect on TLS *in vitro* and in cells. This model is consistent with prior observations that both Pol III core and Pol IV can bind the clamp simultaneously – the toolbelt model (Indiani et al., 2005; Kath et al., 2014; Pagès and Fuchs, 2002). Maintaining the α-cleft interaction is essential to prevent other factors from accessing the P/T junction and compromising processive replication (Dohrmann and McHenry, 2005; Indiani et al., 2009). We propose that during exchange of Pol III holoenzyme, another Pol III core rapidly binds the β_2_ clamp due to the pre-association of Pol III holoenzyme with the replisome through its interaction with the DnaB helicase (Downey and McHenry, 2010; Georgescu et al., 2012) (Reyes-Lamothe et al., 2010) (Figure S6A). In essence this is a “dynamic toolbelt” model, in which rapid exchange of Pol III core happens through the α-cleft while Pol IV carries out synthesis (Figure S6A).

It is likely that the ε-cleft interaction also plays a role in limiting the size of the TLS patch. Pol IV is expected to synthesize only a short patch before it falls off the template and loses its CBM-cleft interaction (transitions from c through f in Figure 6A) (Kath et al., 2014; Wagner et al., 2000). Following the dissociation of Pol IV from the ε-cleft, Pol III core must re-establish its ε-cleft interaction before it can resume processive replication (transition from f to a in Figure 6A). This step would be much faster than expected for diffusion-controlled binding of Pol III core if Pol III remains bound to the β_2_ clamp while Pol IV performs TLS. Conversely, weakening the ε-cleft interaction could result in longer TLS patches and slower replication due to rebinding of Pol IV (Figure 6A, transition from f to d rather than to a).

### Repriming is a failsafe mechanism for rescuing stalled replisomes

We demonstrated that TLS and repriming compete kinetically to resolve stalled replisomes. Failure to carry out timely TLS at the fork results in repriming, which acts as a failsafe to ensure replisome progression. A consequence of repriming is the formation of a ssDNA gap behind the fork (Figure S6B). This gap is initially coated by SSB and subsequently converted to a RecA-ssDNA filament, which induces the SOS DNA damage response. Alternatively, TLS at the fork yields a continuous DNA product and therefore does not lead to induction of the SOS response.

Within our reconstitution, repriming downstream of a N^2^-FFdG adduct on the leading strand template occurred quite slowly compared to Pol IV-mediated TLS at the fork. Similar to prior observations of repriming downstream of CPD lesions, we observed repriming occurring on the timescale of ∼4 minutes, implying that the rate of repriming is likely independent of lesion identity (Yeeles and Marians, 2013; 2011). While direct measurements of the rate of repriming in cells have not been made, estimates (∼10 seconds) suggest that it could be much faster than in vitro observations (Rupp and Howard-Flanders, 1968; Rupp et al., 1971). Therefore, it is possible that other factors present in the cellular environment, but not in current biochemical reconstitutions, may facilitate repriming.

On the other hand, the rate of TLS is controlled by at least two factors: (1) the rate at which a TLS polymerase can bind the P/T junction and (2) the rate at which the TLS polymerase can insert and extend past the lesion. As we have observed here, access of a TLS polymerase to the P/T junction at the replication fork is regulated, in part, by the dynamics of the *ε* gate. Increasing concentrations of Pol IV led to more frequent TLS at the fork within our biochemical reconstitution. Similarly, when the level of Pol IV was constitutively elevated as a result of eliminating LexA repressor binding sites in the promoter of the *dinB* gene (*dinBp-lexA-mut*), the MMS-induced SOS response was decreased relative to the wild-type strain, indicating that creation of ssDNA gaps is suppressed (Figures 6B, S6B and S6C). These results show that higher levels of Pol IV help to overcome the gatekeeping barrier of the ε subunit by enabling Pol IV to more readily capture the *ε* open state.

As cellular concentrations of all three TLS DNA polymerases are increased upon induction of the DNA damage-dependent SOS response, it is likely that TLS at the fork plays a more prominent role in rescuing lesion stalled replisomes after SOS induction. During the early stages of the SOS response, when levels of TLS polymerases are relatively low, the majority of lesion-stalled replisomes are resolved by lesion skipping and most of the resulting ssDNA gaps are repaired by DA (Figure 6B). This is supported by our *in vivo* observations that only small fractions of lesion-stalled replisomes were resolved by TLS, which is likely through a gap-filling process (Figures 5B-5D). A potential exception is Pol IV, which is present at relatively high levels even in the SOS uninduced state, and can efficiently bypass cognate lesions (Figures 1D and 1E). In a *ΔdinB* strain, replisomes stalled at Pol IV cognate DNA lesions would be predominantly resolved by lesion skipping, leaving more ssDNA gaps than in a *dinB*^*+*^ strain. Indeed, when Pol IV-mediated TLS was completely abolished by the deletion of *dinB*, the MMS-induced SOS response was highly elevated compared with the wild type strain (Figure S6C), consistent with an increase in repriming. Collectively, then, pathway choice is a net result of two counteracting molecular events, 1) gatekeeping by the ε subunit and 2) mass action of TLS polymerases (Naiman et al., 2014), with the latter changing throughout the SOS-response. Failure of the TLS polymerase to gain access to the P/T junction or the inability to quickly bypass the lesion upon binding results in repriming and subsequent gap repair.

### The competition between TLS at the fork and repriming and its impact on mutagenesis

TLS at the fork prevents the creation of ssDNA gaps by outcompeting repriming. This gap-suppressing effect of TLS at the fork may play a pivotal role in determining the extent of damage-induced mutagenesis. A recent study has demonstrated that damage-induced mutagenesis is not limited to the lesion site but instead can spread a few hundred nucleotides from the lesion (Isogawa et al., 2018). This extended patch of low-fidelity synthesis likely results from gap filling synthesis where bypass of the lesion by the cognate TLS polymerase is followed by the sequential action of multiple polymerases, including the highly mutagenic Pol V, to fill in the remaining ssDNA gap (Figure 6B) (Fujii et al., 1999; Tang et al., 2000). However, if TLS occurs at the fork, error-prone TLS may be limited to around the blocking lesion, as Pol III can readily regain control of the P/T junction (Figure 6B).

Within cells the extent to which TLS occurs at the fork versus in a ssDNA gap created by repriming remains unclear. Lesion identity, which plays a role in determining the rate of TLS, almost certainly affects the partitioning between these pathways. Lesions that are strongly blocking, such as UV-induced lesions, are likely to be predominantly resolved through repriming due to the absence of Pol V in the early stage of the UV-induced SOS response (Goodman et al., 2016). Even in the presence of highly elevated levels of Pol V, gap filling may still be the predominant mechanism as Pol V activation requires a RecA-ssDNA filament downstream of the stalled replisome (Fujii and Fuchs, 2009; Fujii et al., 2004; Jiang et al., 2009). Weakening the ε gate still dramatically increased Pol V-mediated TLS in cells (Figure 5C), suggesting that the interaction between Pol III and the β_2_ clamp likely influences polymerase switching independent of whether TLS occurs at the fork or in a gap (Figure 6B). Biochemical studies presented here and in prior work, demonstrate that Pol IV can efficiently carry out TLS at the fork for at least a subset of lesions (Gabbai et al., 2014; Ikeda et al., 2014). Cellular evidence for Pol IV-mediated TLS at the fork remains limited but the role of *dinB* in MMS-induced mutagenesis provides circumstantial evidence. 3meA is the major replication-blocking lesion created in MMS-treated cells and is bypassed primarily by Pol IV in a largely error-free manner (Bjedov et al., 2007; Cafarelli et al., 2013; Fu et al., 2012; Larson et al., 1985). Intriguingly, deletion of *dinB* increases MMS-induced mutagenesis, which may result from more mutagenic TLS over 3meA by other polymerases at the fork (Bjedov et al., 2007; Scotland et al., 2015). However, we speculate that this increase is likely related to the creation of ssDNA gaps due to frequent repriming. In the absence of Pol IV, other polymerases mediate TLS over methyl lesions primarily in a post-replicative manner because inefficient synthesis past the lesion is likely outcompeted by repriming (Figure 6B). As is the case for untargeted mutagenesis by UV lesions (Isogawa et al., 2018), the remainder of the gap can be filled in by mutagenic gap-filling synthesis with Pol V being the major mutator. In contrast, MMS-induced mutagenesis is reduced by constitutive expression of Pol IV (Scotland et al., 2015). These observations are consistent with Pol IV-mediated TLS at the fork and suppression of ssDNA gaps (Figure 6B).

Our result that the ε gate antagonizes TLS raises the question if other factors may stimulate TLS in cells and therefore contribute to mutagenesis. Intriguingly, we previously showed that Pol IV is highly enriched near replisomes upon DNA damage in a manner that was only partially dependent on interactions with the β_2_ clamp (Thrall et al., 2017), suggesting that additional interactions, possibly with replisome components, are required for localization of Pol IV to stalled replisomes. Identifying these putative Pol IV– replisome interactions, along with factors that may promote repriming, will be important areas of future investigation.

## Acknowledgements

We thank Shingo Fujii (CNRS, Marseille), Daniel Semlow (Harvard Medical School), Johannes Walter (Harvard Medical School) and members of the Loparo lab for helpful discussions and comments on the manuscript and Deyu Li (University of Rhode Island) and Yinsheng Wang (University of California, Riverside) for providing the N^2^-FFdG- and N^2^-CEdG-containing oligomers, respectively. This work was supported by National Institutes of Health grants R01 GM114065 (to J.J.L), F32 GM113516 (to E.S.T), Agence Nationale de la Recherche (ForkRepair ANR 11 BSV8 017 01 to R.P.F) and the Australian Research Council (DP150100956 and DP180100858 to N.E.D).

## Author Contributions

S.C., K.N., R.P.F and J.J.L designed the experiments. S.C. performed all of the biochemical experiments, cell sensitivity assays and analyzed the resulting data. K.N. performed the *in* vivo TLS assays. Imaging experiments in *E. coli* were carried out by S.C and E.S.T. Recombinant proteins were purified by S.C. and S.J. Bacterial strains were constructed by S.C., K.N. and J.E.K. All authors contributed to the writing and/or editing of the manuscript.

---

**Supplementary figure 1.**
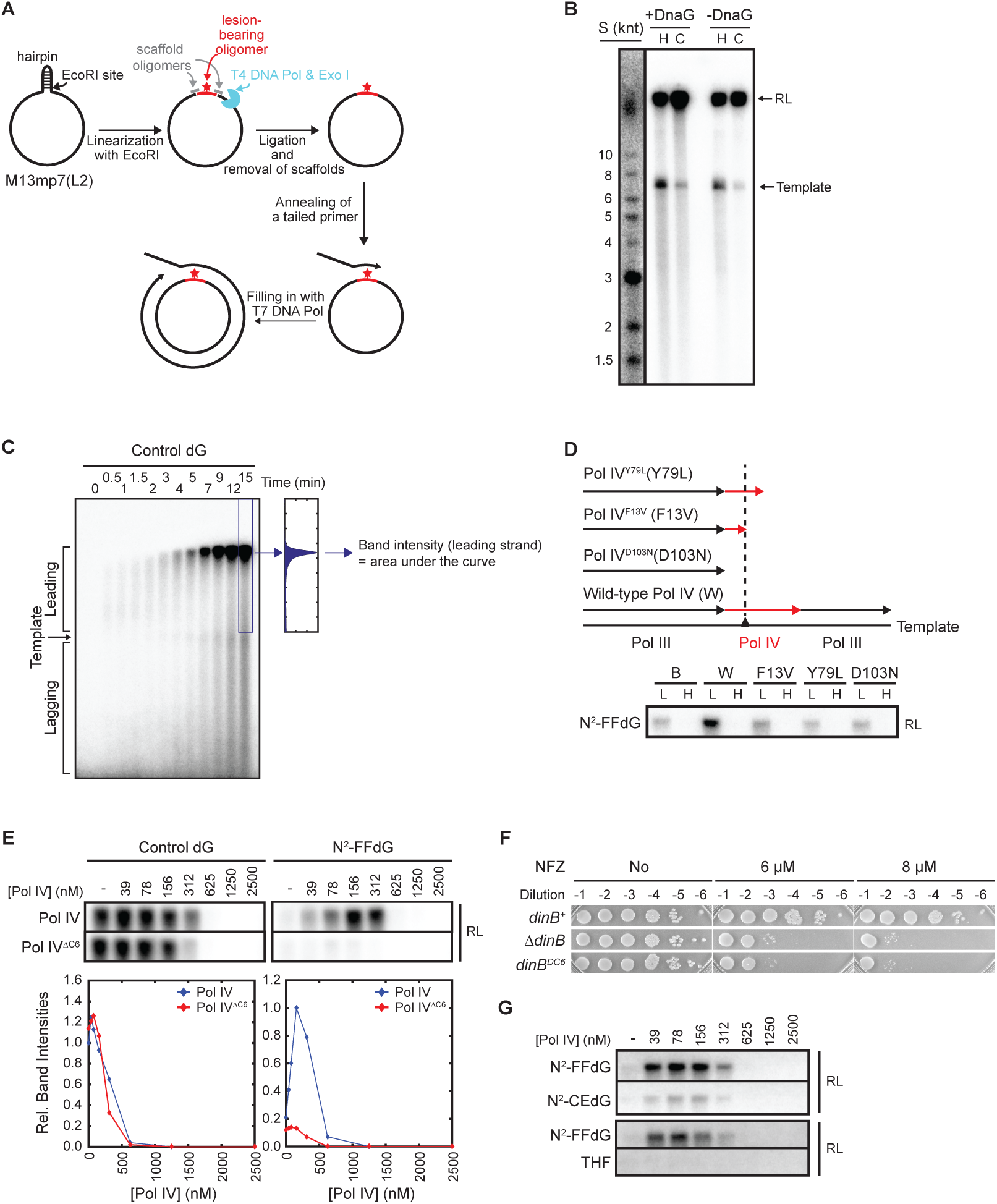
A. An overview of the construction of the N^2^-furfuryl dG (N^2^-FFdG)-containing rolling circle replication template. A single N^2^-FFdG was inserted into a specific position in the leading strand template as described in the methods section. N^2^-CEdG- or THF-containing templates were created using the same procedure (see “Materials and Methods” for details). B. Identification of replication products. A band between the 6 and 8 kilonucleotide (knt) markers (see lane S, ^32^P-end labeled size markers) is unreplicated template. This band resulted from further filling-in of incompletely filled-in rolling circle DNA templates (Wu et al., 1992a; 1992b) and became faint when [α-^32^P]-dATP was added 2 min after the initiation of replication (cold start, C). H refers to hot start in which [α-^32^P]-dATP was added during replication initiation. Smears below the template band are lagging strands and they disappeared in the absence of DnaG. C. Quantitation of leading strand replication products. The same gel as in Figure 1E is shown here as an example. D. Catalytic activity of Pol IV is required for Pol IV-promoted replication of the N^2^-FFdG-containing template but not for inhibiting replication at high concentrations. Only resolution-limited (RL) leading strand products are shown. B, buffer; W, wild-type Pol IV; F13V, Pol IV^F13V^; Y79L, Pol IV^Y79L^; D103N, Pol IV^D103N^; L, 100 nM; H, 1 μM. Pol IV^F13V^ and Pol IV^Y79L^ retain polymerase activities on lesion-free templates but are defective in mediating TLS over N^2^-FFdG (Jarosz et al., 2009). E. Cleft binding activity is absolutely required for Pol IV-mediated TLS over N^2^-FFdG but not for its replication inhibitory activity on the lesion-free control template. (Top) Resolution-limited (RT) leading strand replication products. The concentration range of Pol IV was chosen to cover reported cellular levels of Pol IV under various induction conditions. Pol IV, wild-type full length Pol IV; Pol IV^ΔC6^, Pol IV lacking the cleft binding motif (amino acid 346 to 351 of Pol IV). (Bottom) Integrated intensities of the leading strand replication products. Relative intensities to replication in the absence of Pol IV (for the control template) or in the presence of 156 nM Pol IV (for the N^2^-FFdG-containing template). F. Sensitivity to NFZ showing that Pol IV^ΔC6^ is completely defective in mediating TLS in cells. G. Pol IV mediates TLS over an N^2^-CEdG lesion and THF at the fork less efficiently than it does over N^2^-FFdG.

**Supplementary figure 2.**
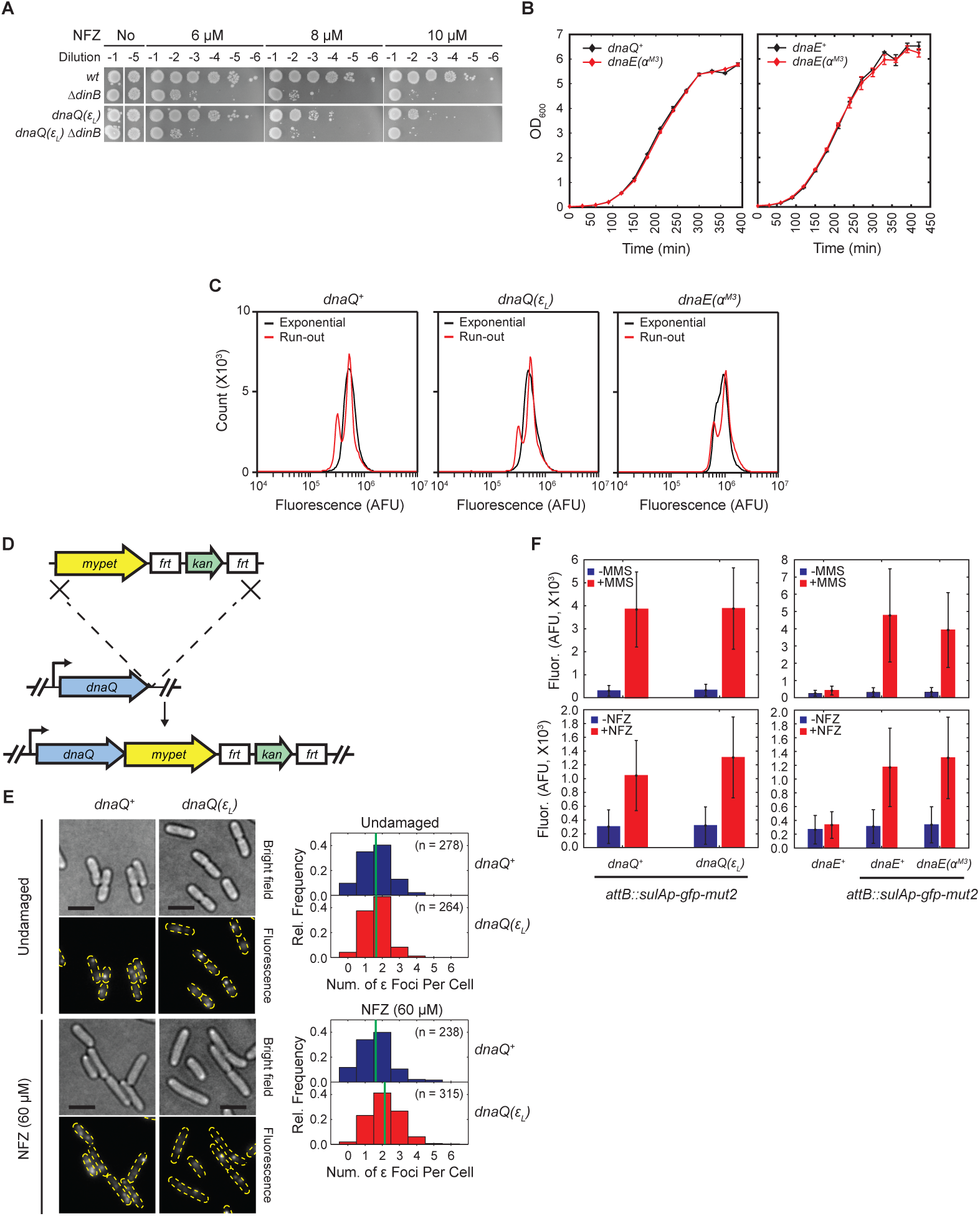
A. Sensitivity assay to NFZ demonstrating that *dnaQ(ε*_*L*_) is epistatic to *ΔdinB*. B. The *dnaQ(ε*_*L*_) and *dnaE(α*^*M3*^) strains retain wild-type growth. Cells were grown in LB with aeration at 37 C. OD_600_ was measured at indicated times. C. *dnaQ(ε*_*L*_) and *dnaE(α*^*M3*^) strains retain wild-type DNA contents. Exponential, asynchronously growing cells in the exponential phase; Run-out, cell division and initiation of a new round of replication in cells growing in the exponential phase were inhibited by cephalexin and rifampicin, respectively. D. Construction of imaging strains expressing either ε-mYpet or ε_L_-mYpet. *mYpet-frt-kan-frt* was appended to the 3’ end of the native *dnaQ* through a linker, in either wild-type or the *dnaQ(ε*_*L*_) strains, by lambda Red recombinase-mediated allelic exchange. E. Integrity of replisomes is not affected by the *dnaQ(ε*_*L*_) mutation. (Left) Representative brightfield and fluorescence micrographs of cells expressing ε-mYPet fusions with cell outlines overlaid on the fluorescence micrographs. (Scale bars: 3 *µ*m.) (Right) Distributions of the number of ε foci per cell. The mean of each distribution is indicated by a green line. F. *dnaQ(ε*_*L*_) and *dnaE(*^*M3*^) strains retain nearly wild-type DNA damage responses. SOS reporter strains (*attB::sulAp-gfp-mut2*) bearing *dnaQ*^*+*^, *dnaQ(ε*_*L*_) or *dnaE(*^*M3*^) alleles were treated with either NFZ (60 µM) or MMS (7 mM) for 1 hour, and fluoresence of GFP-mut2 from individual cells was measured by flow cytometry. Fluor, GFP fluorescence; AFU, arbitrary fluorescence unit.

**Supplementary figure 3.**
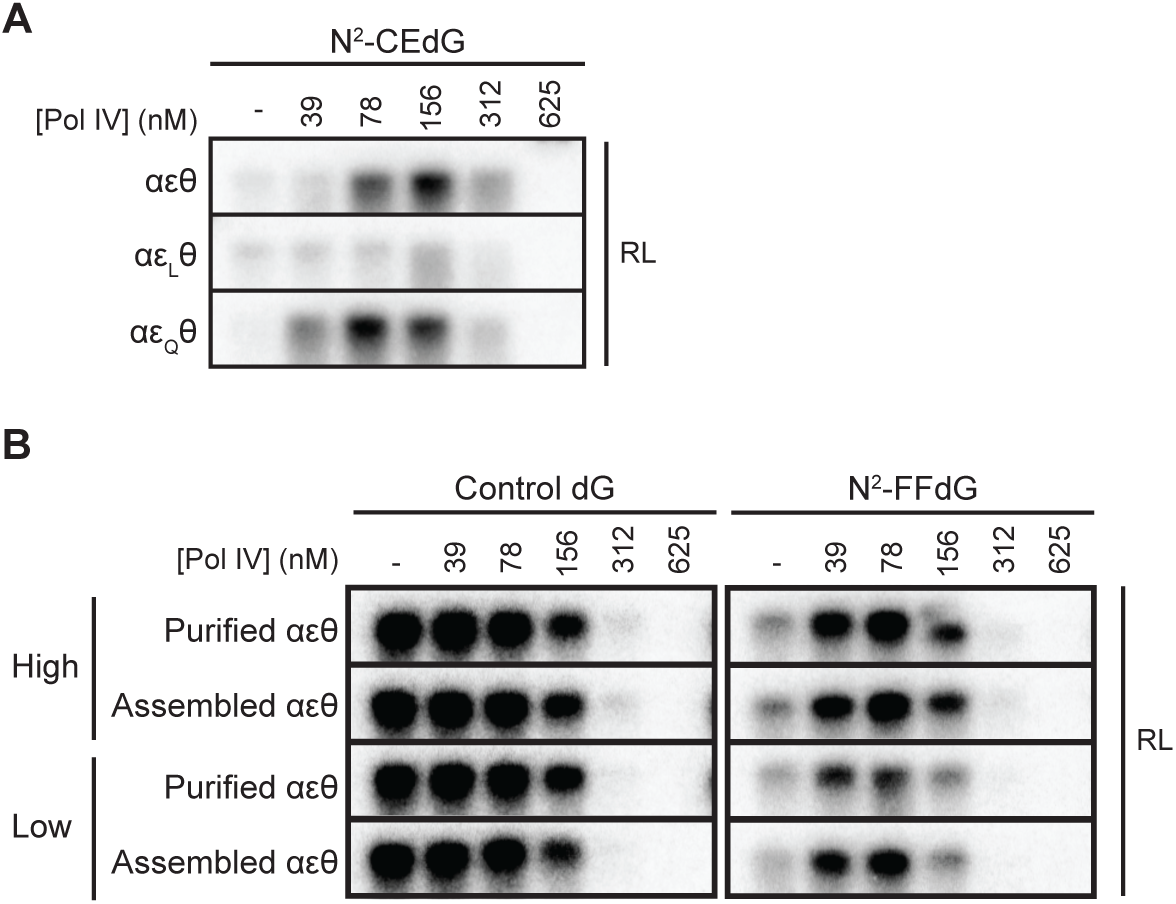
A. Strengthening the ε-cleft interaction (αε_L_θ) suppresses Pol IV-mediated TLS over a N^2^-CEdG lesion. Weakening the interaction (αε_Q_θ) promotes TLS with maximal TLS occurring at lower concentrations of Pol IV as compared to the wild-type replisome (αεθ). Resolution limited (RL) leading strand replication products resulting from the replication of the N^2-^CEdG-containing template by reconstituted replisomes containing the indicated Pol III core complexes. αεθ, wild-type ε-containing replisome; αε_L_θ, ε_L_-containing replisome; αε_Q_θ, ε_Q_-containing replisome. B. Assembled Pol III core complexes retain activities comparable to those of purified Pol III core complexes. Resolution limited (RL) leading strand replication products resulting from the replication of a lesion-free control template or a N^2^-FFdG-containing template by reconstituted replisomes containing either purified or assembled Pol III core complexes (see “Materials and Methods”). High, Pol III core complex: *τ*_3_ complex =4:1; Low, Pol III core complex:_3_ complex=1:1.

**Supplementary figure 4.**
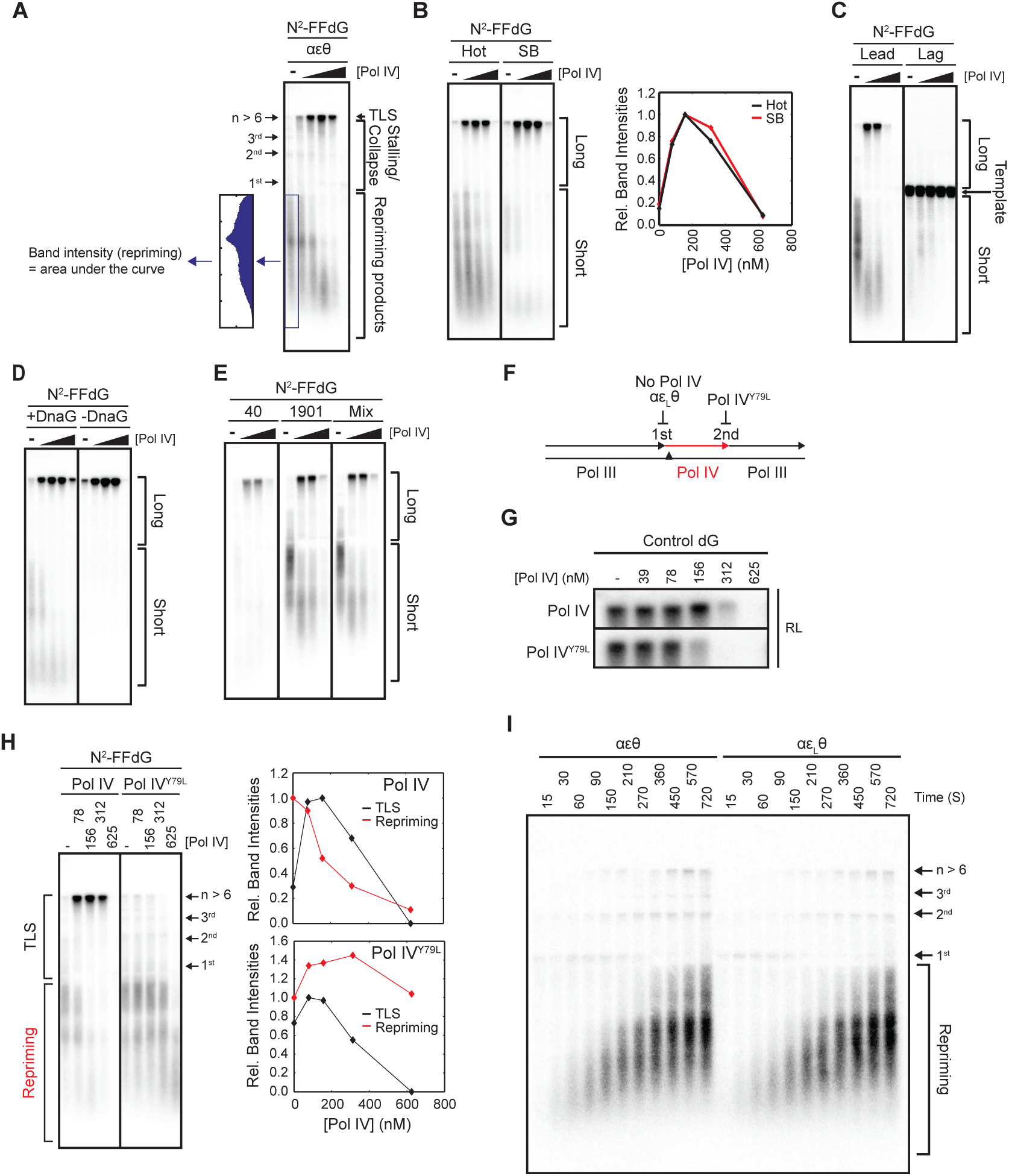
A. Quantitation of repriming. Signals below the template were integrated. The same gel as shown in Figure 4A is reproduced here as an example. B. Detection of replication products of the wild-type replisome (αεθ) by Southern blot with leading strand specific probes: comparison between hot reactions with [α-^32^P]-dATP and Southern blot. (Left) The N^2^-FFdG-containing template was replicated in the presence of varying concentrations of Pol IV (0/78/156/312/625 nM). Replication products (long and short) were labeled with incorporated [^32^P]-dATP during the reaction (Hot) or with ^32^P-labeled oligonucleotides by hybridization after the reaction (SB, Southern blot). Southern blot with leading strand probes detects only a subpopulation of short replication products. (Right) Quantitation of long leading strand replication products. Long, replication products longer than the template; Short, replication products shorter than the template. C. Short leading strand replication products are detected only with leading strand Southern blot probes (Figure 4A). Leading strand probes (Lead) detect both resolution-limited long leading strand products and short leading strand products. Lagging strand probes (Lag) detect templates and short replication products that are distinct from the ones detected with leading strand probes and are presumably Okazaki fragments. D. Short leading strand replication products (<7 knt) are produced only in the presence of DnaG. The N^2^-FFdG-containing template was replicated in the absence (-DnaG) or the presence of DnaG (+DnaG). Replication products were separated in a denaturing agarose gel and detected with leading strand specific Southern blot probes (Figure 4A). E. Short leading strand replication products were more efficiently detected with distal leading strand probes (1901 nts) than with a proximal leading strand probe (40 nts) (Figure 4A). Mix, mixture of two leading strand probes (900 and 1901 nts). F. Steps in polymerase switching during TLS, and the inhibition of each step by indicated genetic modifications (Jarosz et al., 2009; Jergic et al., 2013)}. Pol IV^Y79L^ mediates TLS over the blocking lesion but becomes stalled after incorporating only a few nucleotides past the lesion, preventing second polymerase switching to Pol III. G. Wild-type Pol IV and Pol IV^Y79L^ inhibit replication of a lesion-free template with similar potencies. Resolution-limited (RL) replication products resulting from the replication of a lesion-free control template by the reconstituted replisome in the presence of increasing concentrations of either Pol VI or Pol IV^Y79L^; here the replication products were labeled with [α-^32^P]-dATP. H. Prolonged stalling of replication results in repriming. Addition of Pol IV^Y79L^, a TLS-defective mutant (Figure S4F), which can inhibit replication as potently as the wild-type Pol IV (Figure S4G), led to persistent production of repriming products. This demonstrates that the reduction in repriming in the presence of increasing concentrations of the wild-type Pol IV is due to reduced repriming rather than due to the inhibition of synthesis downstream of the replication block. (Left) Replication of the N^2^-FFdG-containing template in the presence of either Pol IV or Pol IV^Y79L^. Replication products were separated in a denaturing agarose gel and detected by Southern blot. (Right) Quantitation of TLS and repriming products. Amounts of TLS products were plotted relative to the peak amount of TLS products in the presence of Pol IV. Amounts of repriming products were plotted relative to the amount of repriming products in the absence of Pol IV. I. Lesion-stalled wild-type replisomes (αεθ) and ε_L_-containing replisomes (αε_L_θ) reprime downstream of N^2^-FFdG with similar kinetics. The N^2^-FFdG containing template was replicated by either the wild-type or ε_L_-containing replisomes. Replication reactions were quenched after the indicated time of incubation, and replication products were separated in a denaturing gel and detected by Southern blotting with a leading strand probe.

**Supplementary figure 5.**
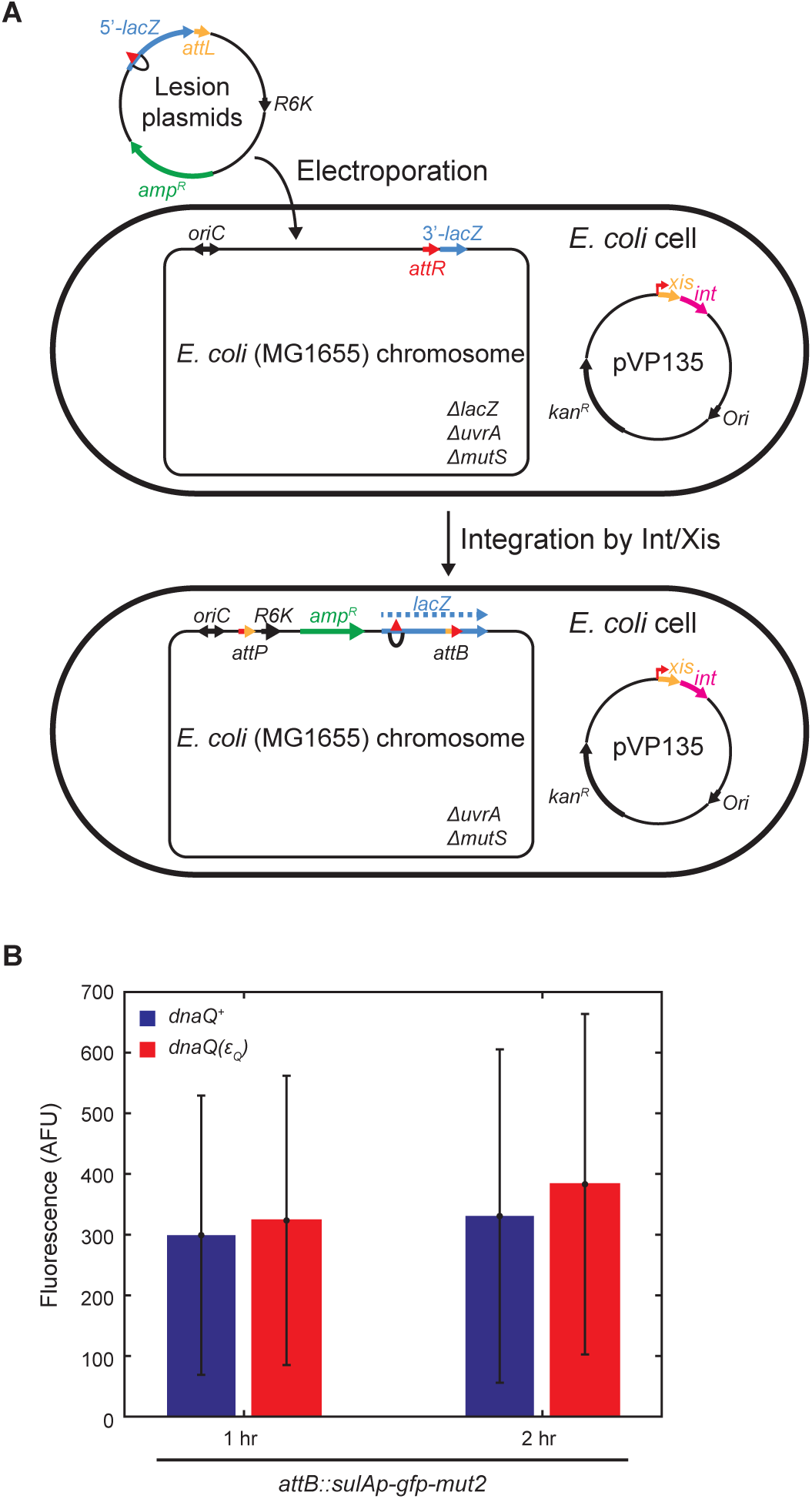
A. Site-specific incorporation of a single DNA lesion into the *E. coli* chromosome. Lesion-containing plasmids are electroporated into recipient strains. Recipient strains express lambda integrase (pVP135) that mediates integration of lesion-containing plasmids into the chromosome at the unique *attR* site. Recipient strains with various *dnaQ* and *umuDC* alleles are described in the main text. B. Basal SOS response levels are not significantly elevated in the *dnaQ(εQ)* strain. Expression levels of GFP-mut2 in mock-treated SOS-reporter strains (*attB::sulAp-gfp-mut2*) bearing either *dnaQ*^*+*^ or *dnaQ(ε*_*Q*_) were mock-treated for 1 or 2 hrs, and fluorescence of GFP-mut2 from individual cells was measured by flow cytometry.

**Supplementary figure 6.**
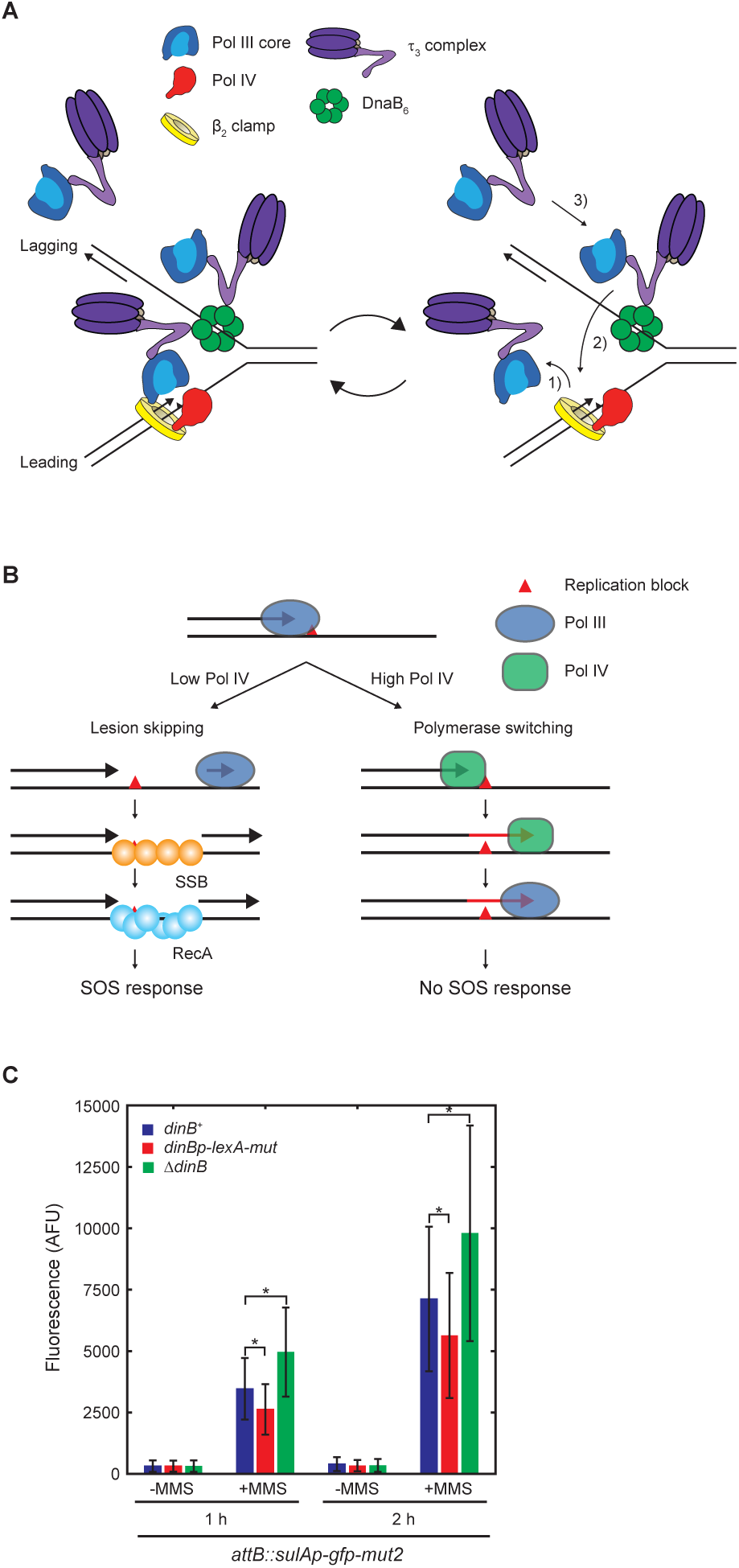
A. Exchange of Pol III holoenzymes during Pol IV-mediated TLS at the fork. 1) Dissociation of Pol III holoenzyme from the stalled replisome. 2) Rapid binding of a Pol III core in a pre-associated Pol III holoenzyme to the α-cleft on the β^2^ clamp at the leading strand lesion. 3) Binding of Pol III holoenzymes in cytosol to the DnaB_6_ helicase at the fork (pre-association with a replisome). B. Induction of the SOS-response by ssDNA gaps generated by repriming downstream of a replication block on the leading strand template (Kowalczykowski et al., 1987; Kowalczykowski and Krupp, 1987; Pagès and Fuchs, 2003; Sassanfar and Roberts, 1990). C. Elevation of expression levels of Pol IV reduces MMS-induced SOS response. SOS reporter strains (*attB::sulAp-gfp-mut2*) bearing the indicated *dinB* loci were treated with MMS (7 mM) for 1 or 2 hours, and fluorescence of GFP-mut2 from individual cells was measured by flow cytometry. *dinB*^*+*^, wild-type dinB locus; *dinBp-lexA-mut*, LexA binding sites in the promoter of *dinB* are mutated; *ΔdinB*, the entire *dinB* gene is replaced with *frt-kan-frt*. *, *p* < 0.0001.

**Supplementary table 1.**
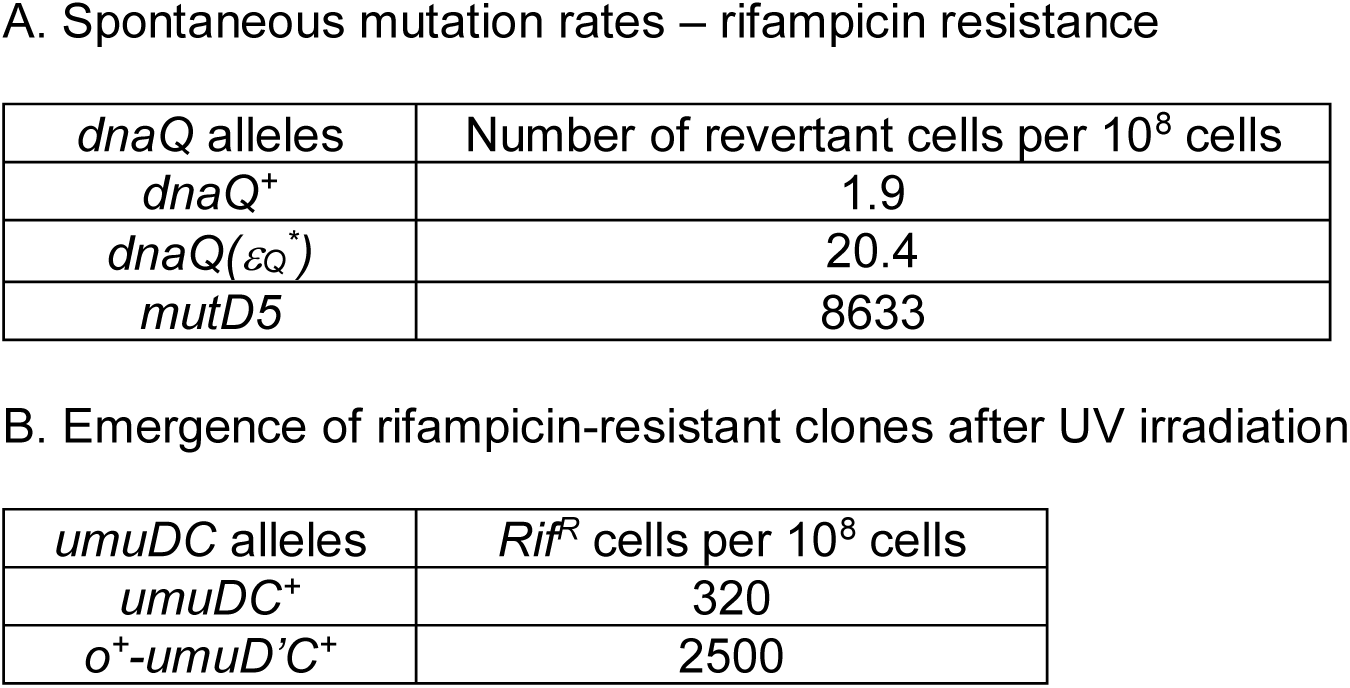
Spontaneous and UV-induced mutation rates.

**Supplementary table 2.**
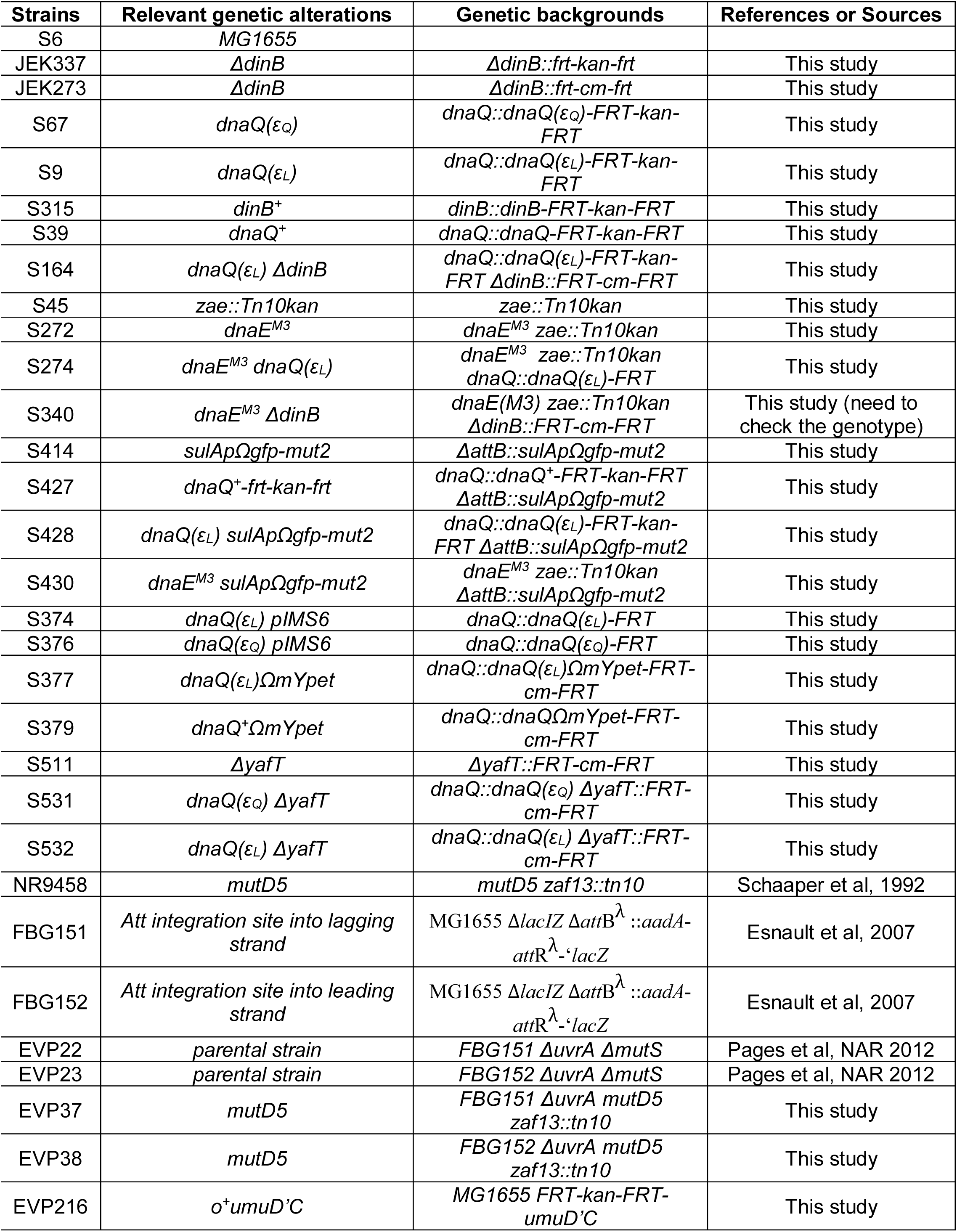
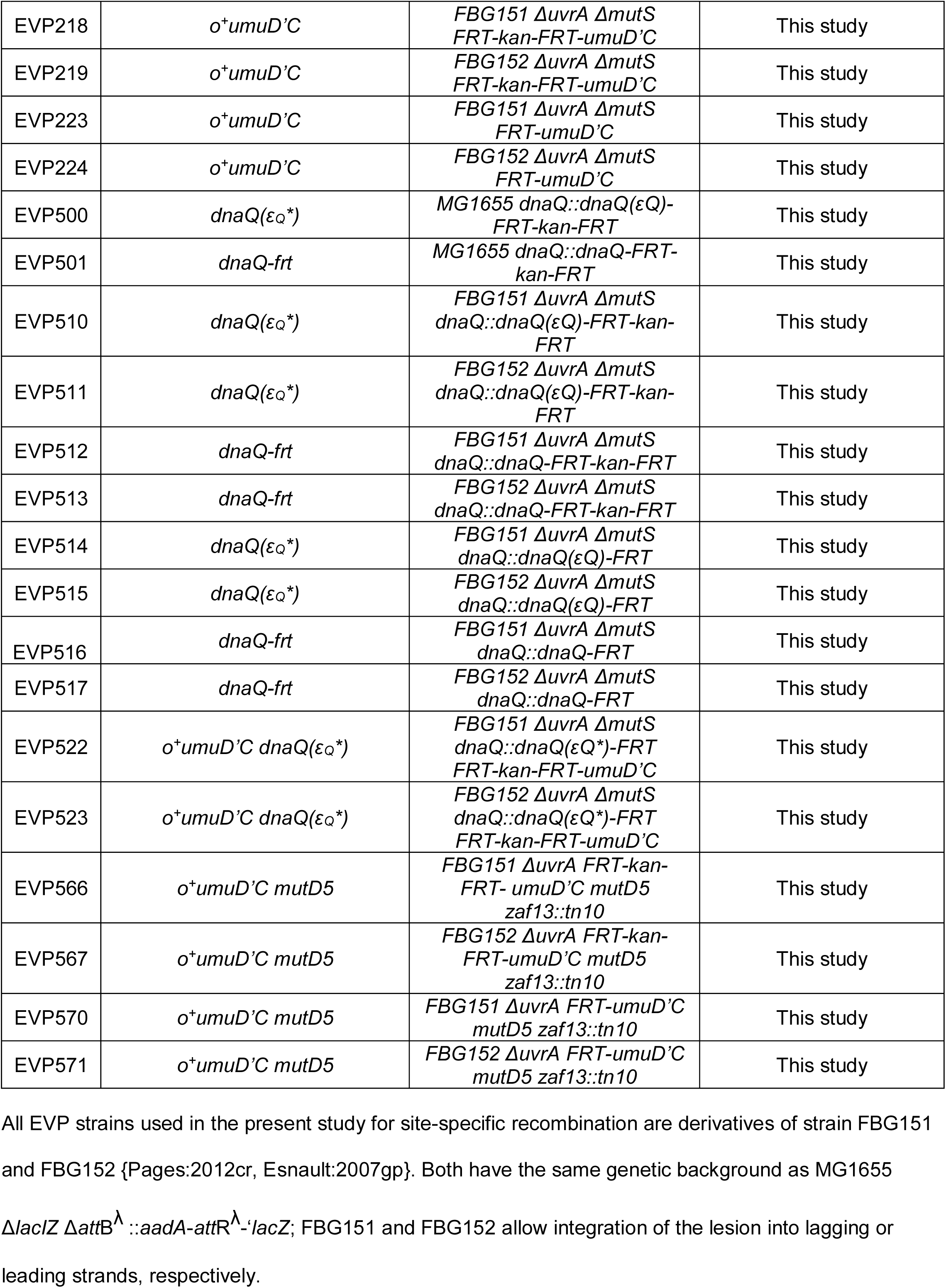

| Strains | Relevant genetic alterations | Genetic backgrounds | References or Sources |
| --- | --- | --- | --- |
| S6 | MG1655 |  |  |
| JEK337 | ΔdinB | ΔdinB::frt-kan-frt | This study |
| JEK273 | ΔdinB | ΔdinB::frt-cm-frt | This study |
| S67 | *dnaQ*($\epsilon_Q$) | *dnaQ*::*dnaQ*($\epsilon_Q$)-FRT-kan-FRT | This study |
| S9 | *dnaQ*($\epsilon_L$) | *dnaQ*::*dnaQ*($\epsilon_L$)-FRT-kan-FRT | This study |
| S315 | *dinB*<sup>+</sup> | *dinB*::*dinB*-FRT-kan-FRT | This study |
| S39 | *dnaQ*<sup>+</sup> | *dnaQ*::*dnaQ*-FRT-kan-FRT | This study |
| S164 | *dnaQ*($\epsilon_L$) ΔdinB | *dnaQ*::*dnaQ*($\epsilon_L$)-FRT-kan-FRT ΔdinB::FRT-cm-FRT | This study |
| S45 | *zae*::Tn10kan | *zae*::Tn10kan | This study |
| S272 | *dnaE*<sup>M3</sup> | *dnaE*<sup>M3</sup> *zae*::Tn10kan | This study |
| S274 | *dnaE*<sup>M3</sup> *dnaQ*($\epsilon_L$) | *dnaE*<sup>M3</sup> *zae*::Tn10kan<br>*dnaQ*::*dnaQ*($\epsilon_L$)-FRT | This study |
| S340 | *dnaE*<sup>M3</sup> ΔdinB | *dnaE*(M3) *zae*::Tn10kan<br>ΔdinB::FRT-cm-FRT | This study (need to check the genotype) |
| S414 | *sulAp*Ωgfp-mut2 | ΔattB::*sulAp*Ωgfp-mut2 | This study |
| S427 | *dnaQ*<sup>+</sup>-frt-kan-frt | *dnaQ*::*dnaQ*<sup>+</sup>-FRT-kan-FRT<br>ΔattB::*sulAp*Ωgfp-mut2 | This study |
| S428 | *dnaQ*($\epsilon_L$) *sulAp*Ωgfp-mut2 | *dnaQ*::*dnaQ*($\epsilon_L$)-FRT-kan-FRT<br>ΔattB::*sulAp*Ωgfp-mut2 | This study |
| S430 | *dnaE*<sup>M3</sup> *sulAp*Ωgfp-mut2 | *dnaE*<sup>M3</sup> *zae*::Tn10kan<br>ΔattB::*sulAp*Ωgfp-mut2 | This study |
| S374 | *dnaQ*($\epsilon_L$) pIMS6 | *dnaQ*::*dnaQ*($\epsilon_L$)-FRT | This study |
| S376 | *dnaQ*($\epsilon_Q$) pIMS6 | *dnaQ*::*dnaQ*($\epsilon_Q$)-FRT | This study |
| S377 | *dnaQ*($\epsilon_L$)ΩmYpet | *dnaQ*::*dnaQ*($\epsilon_L$)ΩmYpet-FRT-cm-FRT | This study |
| S379 | *dnaQ*<sup>+</sup>ΩmYpet | *dnaQ*::*dnaQ*<sup>+</sup>ΩmYpet-FRT-cm-FRT | This study |
| S511 | ΔyafT | ΔyafT::FRT-cm-FRT | This study |
| S531 | *dnaQ*($\epsilon_Q$) ΔyafT | *dnaQ*::*dnaQ*($\epsilon_Q$) ΔyafT::FRT-cm-FRT | This study |
| S532 | *dnaQ*($\epsilon_L$) ΔyafT | *dnaQ*::*dnaQ*($\epsilon_L$) ΔyafT::FRT-cm-FRT | This study |
| NR9458 | *mutD5* | *mutD5* *zaf13*::tn10 | Schaaper et al, 1992 |
| FBG151 | Att integration site into lagging strand | MG1655 ΔlacIZ ΔattB<sup>λ</sup>::*aadA*-attR<sup>λ</sup>-*lacZ* | Esnault et al, 2007 |
| FBG152 | Att integration site into leading strand | MG1655 ΔlacIZ ΔattB<sup>λ</sup>::*aadA*-attR<sup>λ</sup>-*lacZ* | Esnault et al, 2007 |
| EVP22 | parental strain | FBG151 ΔuvrA ΔmutS | Pages et al, NAR 2012 |
| EVP23 | parental strain | FBG152 ΔuvrA ΔmutS | Pages et al, NAR 2012 |
| EVP37 | *mutD5* | FBG151 ΔuvrA *mutD5* *zaf13*::tn10 | This study |
| EVP38 | *mutD5* | FBG152 ΔuvrA *mutD5* *zaf13*::tn10 | This study |
| EVP216 | *o*<sup>+</sup>*umuD*'C | MG1655 FRT-kan-FRT-*umuD*'C | This study |
| EVP218 | *o<sup>+</sup>umuD'C* | *FBG151 ΔuvrA ΔmutS*<br>*FRT-kan-FRT-umuD'C* | This study |
| EVP219 | *o<sup>+</sup>umuD'C* | *FBG152 ΔuvrA ΔmutS*<br>*FRT-kan-FRT-umuD'C* | This study |
| EVP223 | *o<sup>+</sup>umuD'C* | *FBG151 ΔuvrA ΔmutS*<br>*FRT-umuD'C* | This study |
| EVP224 | *o<sup>+</sup>umuD'C* | *FBG152 ΔuvrA ΔmutS*<br>*FRT-umuD'C* | This study |
| EVP500 | *dnaQ(ε<sub>Q</sub><sup>\*</sup>)* | *MG1655 dnaQ::dnaQ(ε<sub>Q</sub>)-*<br>*FRT-kan-FRT* | This study |
| EVP501 | *dnaQ-frt* | *MG1655 dnaQ::dnaQ-FRT-*<br>*kan-FRT* | This study |
| EVP510 | *dnaQ(ε<sub>Q</sub><sup>\*</sup>)* | *FBG151 ΔuvrA ΔmutS*<br>*dnaQ::dnaQ(ε<sub>Q</sub>)-FRT-kan-*<br>*FRT* | This study |
| EVP511 | *dnaQ(ε<sub>Q</sub><sup>\*</sup>)* | *FBG152 ΔuvrA ΔmutS*<br>*dnaQ::dnaQ(ε<sub>Q</sub>)-FRT-kan-*<br>*FRT* | This study |
| EVP512 | *dnaQ-frt* | *FBG151 ΔuvrA ΔmutS*<br>*dnaQ::dnaQ-FRT-kan-FRT* | This study |
| EVP513 | *dnaQ-frt* | *FBG152 ΔuvrA ΔmutS*<br>*dnaQ::dnaQ-FRT-kan-FRT* | This study |
| EVP514 | *dnaQ(ε<sub>Q</sub><sup>\*</sup>)* | *FBG151 ΔuvrA ΔmutS*<br>*dnaQ::dnaQ(ε<sub>Q</sub>)-FRT* | This study |
| EVP515 | *dnaQ(ε<sub>Q</sub><sup>\*</sup>)* | *FBG152 ΔuvrA ΔmutS*<br>*dnaQ::dnaQ(ε<sub>Q</sub>)-FRT* | This study |
| EVP516 | *dnaQ-frt* | *FBG151 ΔuvrA ΔmutS*<br>*dnaQ::dnaQ-FRT* | This study |
| EVP517 | *dnaQ-frt* | *FBG152 ΔuvrA ΔmutS*<br>*dnaQ::dnaQ-FRT* | This study |
| EVP522 | *o<sup>+</sup>umuD'C dnaQ(ε<sub>Q</sub><sup>\*</sup>)* | *FBG151 ΔuvrA ΔmutS*<br>*dnaQ::dnaQ(ε<sub>Q</sub><sup>\*</sup>)-FRT*<br>*FRT-kan-FRT-umuD'C* | This study |
| EVP523 | *o<sup>+</sup>umuD'C dnaQ(ε<sub>Q</sub><sup>\*</sup>)* | *FBG152 ΔuvrA ΔmutS*<br>*dnaQ::dnaQ(ε<sub>Q</sub><sup>\*</sup>)-FRT*<br>*FRT-kan-FRT-umuD'C* | This study |
| EVP566 | *o<sup>+</sup>umuD'C mutD5* | *FBG151 ΔuvrA FRT-kan-*<br>*FRT- umuD'C mutD5*<br>*zaf13::tn10* | This study |
| EVP567 | *o<sup>+</sup>umuD'C mutD5* | *FBG152 ΔuvrA FRT-kan-*<br>*FRT-umuD'C mutD5*<br>*zaf13::tn10* | This study |
| EVP570 | *o<sup>+</sup>umuD'C mutD5* | *FBG151 ΔuvrA FRT-umuD'C*<br>*mutD5 zaf13::tn10* | This study |
| EVP571 | *o<sup>+</sup>umuD'C mutD5* | *FBG152 ΔuvrA FRT-umuD'C*<br>*mutD5 zaf13::tn10* | This study |

All EVP strains used in the present study for site-specific recombination are derivatives of strain FBG151 and FBG152 {Pages:2012cr, Esnault:2007gp}. Both have the same genetic background as MG1655 $\Delta lacI Z \Delta attB^{\lambda}::aadA-attR^{\lambda}-lacZ$ ; FBG151 and FBG152 allow integration of the lesion into lagging or leading strands, respectively.

**Supplementary table 3.**

| **Plasmid ID** | **Plasmids** | **References or Sources** |
| --- | --- | --- |
| SE258 | pKD3-sulAp-gfp-mut2 | This study |
| JEK541 | pKD4-dnaQ | This study |
| JEK543 | pKD4-dnaQ($\epsilon_L$) | This study |
| SE167 | pMAK700-dnaE<sup>M3</sup> | This study |
| SE100 | pKD4-dinB | This study |
| JEK1 | pET11T-dinB | This study |
| SE157 | pET11T-dinB<sup>Y79L</sup> | This study |
| SE289 | pET11a-his6-ubq-eCTS | This study |
| SE307 | pET-his6-ubq-eCTS($\epsilon_L$) | This study |
| SE228 | pET11a-dnaQ<sup>D12A,E14A</sup> | This study |
| SE229 | pET11a-dnaQ($\epsilon_L$)<sup>D12A,E14A</sup> | This study |
| pGP1 | Description in Materials and Methods section | Pages et al, NAR 2012 |
| pGP2 | Description in Materials and Methods section | Pages et al, NAR 2012 |
| pGP9 | Description in Materials and Methods section | Ikeda et al, NAR 2014 |
| pVP135 | Description in Materials and Methods section | Pages et al, NAR 2012 |
| pVP141 | Description in Materials and Methods section | Pages et al, NAR 2012 |
| pVP142 | Description in Materials and Methods section | Pages et al, NAR 2012 |
| pVP143 | Description in Materials and Methods section | Pages et al, NAR 2012 |
| pVP144 | Description in Materials and Methods section | Pages et al, NAR 2012 |
| pVP146 | Description in Materials and Methods section | Pages et al, NAR 2012 |
| pKN3 | pGB2-FRT-Kan-FRT-umuD'C | This study |
| pKN14 | pMK-dnaQ($\epsilon_Q^*$) - FRT-Kan-FRT | This study |
| pKN15 | pMK-dnaQ - FRT-Kan-FRT | This study |

## Materials and Methods

### Proteins

#### Protein purification

Purified αεθ, αε_Q_θ, αε_L_θ, α, ε, ε_Q_, ε_L_, θ, DnaB, clamp loader complex (τ_3_δδκψ), β_2_ clamp, DnaG and SSB were either purified as previously described (Gulbis et al., 2004; Hamdan et al., 2002; Jergic et al., 2013; Maki and Kornberg, 1988; Ozawa et al., 2005; San Martin et al., 1995; Stamford et al., 1992; Tanner et al., 2008; Wijffels et al., 2004; Williams et al., 2002; Xiao et al., 1993). Wild-type Pol IV and its variants were expressed in BL21(DE3) pLysS lacking the native *dinB* and purified as previously described (Jarosz et al., 2006; Kath et al., 2014). α^M3^ was expressed as a N-terminal fusion of His10-SUMO in BL21(DE3) pLysS and purified by Ni-NTA affinity purification. N-terminal His10-SUMO was cleaved by Ulp1, creating untagged α^M3^, and cleaved His10-SUMO was removed by Ni-NTA affinity purification.

#### Assembly of various polymerase complexes

α_V832G_εθ, αε^D12A,E14A^θ and αε_L_ ^D12A,E14A^θ: To reconstitute these Pol III core complex, purified α subunits, ε subunits and θ subunits were mixed at 1:1.2:1.2 (concentration of the α subunit was 5 αM) ratio and incubated at room temperature for 3 min and the mixtures were further incubated on ice for 30 min before used in *in vitro* replication assays. Pol III core complex assembled in this way behaved in the same way as those assembled and purified by ion-exchange chromatography (Figure S3B).

#### Chemicals

Methyl methanesulfonate (Sigma, 129925), Nitrofurazone (Fluka, PHR1196), PicoGreen (Molecular probes, P7581)

### Rolling circle replication

#### Construction of control and lesion-containing rolling circle DNA templates

To make a lesion containing circular ssDNA, a lesion containing oligomer-1) 5’- CTACCTXTGGACGGCTGCGA-3’ (X= N^2^-FFdG, N^2^-furfuryl dG), 2) GCGCAAAXCTTGAGCTC (X=N^2^- CEdG, (1-carboxyethyl)-2’-deoxyguanosine), 3) 5’-CTACCTXTGGACGGCTGCGA-3’ (X=THF, tetrahydrofuran), 4) 5’-CTACCTXTGGACGGCTGCGA-3’ (X=dG)- was ligated into EcoRI-linearized M13mp7L2 as previously described (Figure S1A) (Delaney and Essigmann, 2006). A lesion free or a lesion-containing circular ssDNA was converted into a 5’-tailed dsDNA by T7 DNA polymerase-catalyzed extension of a primer (T_36_GAATTTCGCAGCCGTCCACAGGTAGCACTGAATCATG) that was annealed over the lesion-containing region of the circular ssDNA (Figure S1A). At each enzymatic step, enzymes were heat-inactivated and removed by standard phenol-chloroform extractions. N^2^-FFdG- and N^2^-CEdG-containing oligomers were kindly provided by Deyu Li (University of Rhode Island) and Yinsheng Wang (University of California, Riverside) respectively. The control and THF-containing oligomers were synthesized by Integrated DNA Technologies (Skokie, IL, USA).

#### Rolling circle replication

Rolling circle replication reactions proceed through the following steps. Replisome assembly at the replication fork: Template (1 nM), τ_3_δδκψ (20 nM), indicated Pol III core (20 nM), β_2_ clamp (20 nM), DnaB_6_ (50 nM), ATPψS (50 μM) and dCTP/dGTP (60 μM each) were mixed in HM buffer (50 mM Hepes (pH 7.9), 12 mM Mg(OAc)2, 0.1 mg/ml BSA and 10 mM DTT) on ice: concentrations in the parentheses were final concentrations in the reactions. Assembly reaction mixtures were then transferred to a water bath at 37 °C and incubated for 4 to 6 min.

Initiation of replication: 10X initiation mixtures of ATP (1 mM), CTP/GTP/UTP (250 μM each), dATP/dTTP (60 μM each), [α-^32^P]-dATP, SSB_4_ (200 μM), DnaG (100 nM) were made and kept on ice; concentrations in the parentheses were final concentrations in the reactions. Replication was initiated by the addition of initiation mixture; 10X initiation mixtures were prewarmed at 37 °C for 1 min before added to the assembly reactions.

Replication reaction: After incubation at 37 °C for 12 min unless indicated otherwise, replication reactions were quenched by adding EDTA (final 25 mM).

Replication products were separated in a 0.6 % alkaline denaturing alkaline agarose gel and visualized by autoradiography. Radioactive signals were quantitated using ImageJ. Synthesis of leading strand was determined by integrating signals above the template (Figure S1C). These values for the lesion-free control template and for N^2^-FFdG-containig template were defined “processive replication” and “TLS” respectively. Relative band intensities were calculated with respect to the band intensity in the absence of Pol IV for the lesion-free template and the maximal band intensity for the lesion containing templates in the presence of Pol IV, respectively. To compare Pol IV-mediated TLS among various Pol III cores, replication of the N^2^- FFdG containing template was first normalized with replication of a lesion-free control template; relative TLS was then calculated using these normalized values. Replication by αεθ-containing replisome was included in all experiments as a reference. In some cases, only resolution-limited bands were presented for easy comparison among various conditions.

#### Detecting repriming products by Southern blot

Rolling circle replication reactions were performed as described above but without [α-^32^P]-dATP. Replication products were separated in a 0.6% denaturing agarose gel and mildly cleaved by incubating the gel in 0.25 M HCl followed by incubating in a denaturing buffer (3 M NaCl/0.4 M NaOH). Cleaved replication products were transferred to a Nylon membrane (Hybond – XL, GE Healthcare) by a downward transfer method with transfer buffer (1.5 M NaCl/0.4 M NaOH). Transferred DNAs were UV cross-linked to the membrane with a UV cross-linker (UV Stratalinker 2400, Stratagene). The cross-linked membranes were incubated in a blocking buffer (ULTRAhyb, Invitrogen, AM8670) for 5 hours. 5’ [^32^P]-labeled oligomer probes (shown in Figure 4A) were added to the blocking buffer and hybridized with the membrane in a hybridization chamber for 20 hours at 42 °C with continuous agitation. Unless otherwise stated, a mixture of leading strand probes (900 and 1901 nts) was used to increase the signals. Radioactive signals were visualized by autoradiography. To quantitate downstream synthesis (repriming), signals below the unreplicated template were integrated using ImageJ as shown in Figure S4A.

### Measuring DNA contents and SOS-response

#### DNA contents

DNA contents were measured with cultures in exponential growth or under a run-out condition. Overnight cultures were diluted to OD_600_∼0.01 and the diluted cultures were further grown at 37 °C with aeration until OD_600_ reached between 0.2 and 0.3. To measure DNA contents of exponentially growing cells, cells were fixed with ethanol (70 % final) and stained with PicoGreen dye for 1 hour in the dark. Fluorescence from individual cells were measured with a flow cytometer (BD Accuri C6).To measure the DNA contents of cells under a run-out condition, cultures were treated with cephalexin (30 μg/ml final) and rifampicin (300 μg/ml final) and further incubated for 3 hours at 37 °C with aeration. Then cells were processed in the same way as for measuring exponentially growing cells (Ferullo et al., 2009). In total 10^5^ cells were measured.

#### SOS-response

Overnight cultures of SOS reporter strains were diluted into Luria broth (LB) to OD_600_ of 0.05 and further cultured at 37 °C with aeration until OD_600_ reached ∼0.3. Cultures were treated with NFZ (60 µM final) or MMS (7 mM final) and further incubated at 37°C with aeration. Aliquots of cultures were removed at varying time points, and cells were fixed with formaldehyde (4 %). Fixed cells were thoroughly washed with phosphate-buffered saline (PBS, pH 7.4) and finally resuspended in PBS. GFP fluorescence from individual cells was measured with a flow cytometer (BD Accuri C6).

#### Measuring sensitivity to NFZ and MMS

Overnight cultures were diluted in 0.9 % NaCl to OD_600_ = 1.0 and serially diluted by 10 fold to 10^6^ fold. Serially diluted cultures were spotted on LB-agar containing indicated concentrations of NFZ or MMS. After 15 hour incubation at 37 °C, plates were photographed.

#### *E. coli* strains

Refer to the supplementary table 2 for *E. coli* strains used in this study

### Construction of strains

#### *dnaQ* strains (S9, S39 and S67)

Strains with different alleles of *dnaQ* were generated using the lambda Red recombinase-mediated homologous alleleic exchange (Sharan et al., 2009). The wildtype allele of *dnaQ* was first subcloned between nucleotide number 30 and 31 of pKD4, upstream of the *frt-kan-frt* cassette, by Gibson assembly (Datsenko and Wanner, 2000; Gibson et al., 2009), generating the plasmid pKD4-*dnaQ*. The cleft-binding motif residues of *dnaQ* (QTSMAF) were mutated by site-directed mutagenesis to generate the plasmid pKD4-*dnaQ(ε*_*Q*_) (CBM residues: ATSMAF) and the plasmid pKD4-*dnaQ(ε*_*L*_) (CBM residues: QLSLPL) (Jergic et al., 2013).

Linear dsDNAs containing the respective *dnaQ* alleles (*ε*_*WT*_, *ε*_*Q*_ or *ε*_*L*_) and the *frt-kan-frt* were amplified from pKD4-*dnaQ*, pKD4-*dnaQ(ε*_*Q*_) and pKD4-*dnaQ(ε*_*L*_) and used to replace native *dnaQ* locus by lambda Red allelic exchange in either the strain MG1655 bearing the plasmid pSIM5 (for *dnaQ*^*+*^ and *dnaQ(ε*_*L*_)) or the strain CH1358 (for *dnaQ(ε*_*Q*_)). CH1358 contains a temperature-inducible genomic copy of the lambda Red operon and a *mutS* mutation for more efficient recombineering, and was a gift of Diarmaid Hughes (Uppsala University). Respective *dnaQ* alleles were verified by sequencing and transferred into fresh isolates of wild-type MG1655 by general P1 transduction.

#### Deletion strains (JEK273, JEK337, and S511)

The entire coding sequences were replaced with *frt-cm*^*R*^*-frt* or *frt-kan*^*R*^*-frt* cassette of pKD3 or 4 respectively by lambda Red allelic exchange in MG1655 bearing pSIM5 as previously described (Sharan et al., 2009); sequences for the starting codon, ultimate 6 amino acids and the stop codon were preserved. Selection markers were flipped out by transforming the cells with pCP20, which expresses a flippase. The removal of a selection marker was selected for sensitivity to the corresponding antibiotics and confirmed by PCR. pCP20, which contains a temperature-sensitive origin of replication, was cured by growing cells at 37 °C.

#### *dnaE^M3^* (S272)

cDNA for *dnaE* was subcloned into pMAK between HindIII and XbaI by Gibson assembly to construct pMAK-*dnaE*. The *dnaE*^*M3*^ mutation (^920^QADMF^924^ to ^920^QLDLF^924^) was introduced to pMAK-dnaE by Quick change method to construct pMAK-dnaE^M3^. S45 (*zae::Tn10kan*) was transformed with pMAK-*dnaE*^*M3*^ and selected for normal growth at 37 °C (Hamilton et al., 1989). Clones were screened for the *dnaE*^*M3*^ mutation by sequencing.

#### dnaE^M3^ dnaQ(ε_L_)

*dnaE*^*M3*^ *zae::Tn10kan* was transferred from S272 into S9 by P1 transduction

#### *sulApΩgfp-mut2* reporter strains (S414, 428 and 429)

*sulAp*, the promoter of the *E. coli* sulA gene (−67 to −1 of *sulA*) was fused to 5’ end of gfp-mut2, and this fusion sequence was subcloned between nucleotide number 30 and 31 of pKD3 by Gibson assembly, generating pKD3-sulAp-gfp-mut2. A linear dsDNA encoding sulAp-gfp-mut2-FRT-cm-FRT was PCR-amplified from pKD3-sulA-gfp-mut2 and used to precisely replace 15 bp *attB* site by red-recombinase-mediated recombineering (S414). Various *dnaQ* alleles were transferred from S39 or S9 into S414 by P1 general transduction to create S427 and S428. *dnaE*^*M3*^ *zae::Tn10kan* was transferred from S272 into S414 by general P1 transduction. Each allele was confirmed by sequencing *dnaQ* or *dnaE* loci.

#### *dnaQ-mypet* strains (S377 and S379)

The *(g4s)X4-mypet-frt-kan-frt* sequence was PCR amplified and introduced to the 3’ end of the *dnaQ* gene by allelic exchange in S374 and S376, creating S377 and S379. pSIM6 was cure by incubating cells at 37 °C.

#### Strains for lesion tolerance monitoring

Strains used for monitoring lesion tolerance were derived from strain FBG151 or FBG152, in which both *mutS* and *uvrA* genes were disrupted (Esnault et al., 2007; Pagès et al., 2012). Strains FBG151 and FBG152 allow integration of a single lesion in the lagging and leading strands, respectively.

#### EVP514, 515, 516 and 517

The *dnaQ(ε*_*Q*_*\*)* mutation (^182^QTSMAF^187^ to ^182^QTSAAA^187^) was introduced into the native *dnaQ* gene by lambda red recombinase-mediated allelic exchange. First the entire *dnaQ* sequence bearing the *dnaQ(ε*_*Q*_*\*)* mutation followed by *frt-kan-frt* (*dnaQ((ε*_*Q*_*\*)-frt-kan-frt*) and the sequence immediate downstream of the *dnaQ* gene (corresponding to +1 to +111 of *dnaQ* gene) was subcloned into pMK at the Sfil site, producing pKN14; pNK14 was created by Life Technologies. We restored the wildtype CBM sequence in pKN14 by site-directed mutagenesis using primers VP264/VP265 (5’-ACCGGTGGTCAAACGTCGatgGCTtttGCGATGGAAGGAGAGACA-3’ and 5’-TGTCTCTCCTTCCATCGCAAAAGCCATCGACGTTTGACCACCGGT-3’), producing pKN15. To replace the native *dnaQ* gene with either *dnaQ(εQ*)-frt-kan-*frt or dnaQ*-frt-kan-frt*, pKN14 and 15 were digested with EcoRV and SspI, creating knocking-in (KI) cassettes for allelic exchange. MG1655 expressing lambda red recombinase was transformed with each of these KI cassettes and selected for *kan*^*R*^, producing EVP500 (MG1655 *dnaQ(εQ*)-frt-kan-*frt) and EVP501 (MG1655 dnaQ*-frt-kan-frt)* respectively. These *dnaQ* alleles were transferred from EVP500/EVP501 into EVP22/23 (FBG151/FBG152 *ΔuvrA ΔmutS)* by P1 transduction, producing EVP510/511 (FBG151/FBG152*ΔuvrA ΔmutS* dnaQ*(εQ*)-frt-kan-frt*), and EVP512/513 (FBG151/FBG152*ΔuvrA ΔmutS* dnaQ*-frt-kan-frt*). The kan cassette in these strains was flipped out by transiently expressing flippase, producing EVP514/515 (FBG151/FBG152*ΔuvrA ΔmutS dnaQ::dnaQ(εQ)-FRT)* and EVP 516/517*(FBG151/FBG152ΔuvrA ΔmutS dnaQ-FRT)*.

#### EVP223 and 224

The *umuD’C* mutation was introduced into the native *umuDC* gene by lambda red recombinase-mediated allelic exchange. For this, we modified pRW134 (a gift from Roger Woodgate), pGB2-based plasmid, which bears *umuD’C* preceded by the native *umuDC* promoter (Ennis et al., 1995); in *umuD’C*, the DNA sequence encoding amino acid 1 to 14 of UmuD was replaced with ATG. To introduce a selection marker, *frt-kan-frt* sequence from pKD13 was subcloned at the PmII site in pRW134, which is located near the 5’ end of the promoter of *umuDC*, producing pKN3 (pGB2-*frt-kan-frt-umuD’C*); note that the native *umuDC* promoter is preserved between *frt-kan-frt* and *umuD’C*. pKN3 was digested with ScaI and EcoRI, producing *frt-kan-frt-umuD’C*. MG1655 expressing lambda recombinase was transformed with frt-kan-frt-umuD’C KI cassette, producing EVP216 (MG1655 *frt-kan-frt-umuD’C*). The umuD’C allele was transferred by P1 transduction from EVP216 to EVP22/23 (FBG151/*FBG152 ΔuvrA ΔmutS*), producing EVP218/219 (FBG151/*FBG152 ΔuvrA ΔmutS frt-kan-frt-umuD’C)*. The kan cassette in these strains was flipped out by transiently expressing flippase, producing EVP223/224 (FBG151/*FBG152 ΔuvrA ΔmutS frt-umuD’C)*

#### EVP522 and 523

*frt-kan-frt-umuD’C* was transferred from EVP216 into EVP514/515 (FBG151/FBG152 *ΔuvrA ΔmutS dnaQ::dnaQ(εQ)-FRT)* by P1 transduction producing EVP522/523 (FBG151/FBG152*ΔuvrA ΔmutS dnaQ::dnaQ(εQ)-FRT frt-kan-frt-umuD’C)*.

#### EVP570 and 573

Strain NR9458 *(mutD5 zaf13:tet)* was a gift from Roel Schaaper (Schaaper and Cornacchio, 1992). The *mutD5* allele was transferred by P1 transduction from NR9458 into EVP20/21 (FBG151/FBG152*ΔuvrA*), producing EVP37/38 (FBG151/FBG152*ΔuvrA mutD5)*. Subsequently, *frt-kan-frt-umuD’C* was transferred by P1 transduction from EVP216 into EVP37/38, producing *EVP566/EVP567* (FBG151/FBG152 *ΔuvrA mutD5 frt-kan-frt-umuD’C)*. The kan cassette in these strains was flipped out by transiently expressing flippase, producing EVP570/573 (FBG151/*FBG152 ΔuvrA frt-umuD’C mutD5)*.

### Live cell imaging of ε foci

#### Sample preparation and NFZ treatment

Single colonies were isolated by streaking glycerol stocks on LB plates containing 30 µg/mL kanamycin and incubating overnight at 37 °C. The day before imaging, a 3 mL LB culture was inoculated with a single colony and incubated in a roller drum at 37 °C for approximately 8 h. A 3 mL culture in M9 medium supplemented with 0.4% glucose, 1 mM thiamine hydrochloride, 0.2% casamino acids, 2 mM MgSO_4_, and 0.1 mM CaCl_2_ was inoculated with a 1:1,000 dilution of the LB culture and incubated overnight under the same conditions. Imaging cultures in 50 mL of supplemented M9 medium were then inoculated with a 1:200 dilution of the overnight M9 culture and incubated at 37 °C shaking at 225 rpm.

Cells were harvested when imaging cultures reached OD_600nm_ ∼ 0.15. An aliquot of 1 mL was removed and concentrated by centrifugation at 8,609 × *g*. Agarose pads were prepared by dissolving GTG agarose (NuSieve) to a 3% concentration in M9 medium supplemented with 0.4% glucose, 2 mM MgSO_4_, and 0.1 mM CaCl_2_ on a heat block at 65 °C. Pads were cast by depositing 500 µL of molten agarose between two 25 × 25 mm coverslips (VWR) that were rinsed with ethanol and deionized (DI) water. A small volume of 1 µL or less of the concentrated culture was pipetted on a piece of the agarose pad and sandwiched between another coverslip, which was cleaned by two alternating 30 min cycles of sonication in ethanol and then 1 M KOH. Coverslips cleaned by sonication were rinsed thoroughly with deionized (DI) water and stored in DI water until needed.

For nitrofurazone (NFZ) treatment, a 60 mM stock solution of NFZ was prepared freshly in dimethylformamide. A 1:1,000 dilution of this stock was added to the imaging cultures at OD_600nm_ ∼ 0.15, resulting in a 60 µM final concentration. Cultures were grown for an additional 1 h and then prepared for imaging as above.

#### Microscopy

Live cell imaging was carried out on a customized Nikon TE2000 microscope equipped with a Nikon CFI Apo 100X/1.49 NA TIRF objective using 514 nm laser excitation (Coherent Sapphire, 150 mW). After expansion by a telescope, the beam passed through an excitation filter (Chroma ZET405/20X, ZET514/10X), a dichroic beam combiner (Chroma ZT514rdc), and a second telescope. Finally, a 400 mM focal length lens was used to focus the beam off-center in the objective back focal plane to achieve highly inclined thin illumination. [1] Imaging was performed using a single-band TIRF filter cube designed for mYPet (Chroma TRF49905, containing a ZT514rdc dichroic filter, a ET545/40m emission filter, and a ET525lp longpass filter) and a Hamamatsu ImageEM C9100-13 EMCCD camera. The final pixel size, with the internal 1.5X microscope magnification, was approximately 106 nm. All movies were recorded using an integration time of 13.283 ms and a 514 nm laser power of approximately 16.6 W cm^−2^, and corresponding brightfield images of cells were recorded using white light transillumination. For each strain and condition, two independent experiments were performed on separate days.

#### Image analysis

Automated image analysis was performed in MATLAB. Bright-field images were analyzed using the MicrobeTracker package (Sliusarenko et al., 2011) to determine the cell outlines. Cells at the edge of the field of view were excluded from analysis. The first 10 frames of fluorescence movies were averaged and analyzed using the package u-track (Jaqaman et al., 2008). Within each cell outline, the u-track point source detection algorithm (Aguet et al., 2013) was used to identify ε-mYPet foci and fit them to symmetrical 2D Gaussian approximations of the point-spread function (PSF). For each focus, the Gaussian x and y centroid, amplitude, width (δ), and background offset were determined. With the exception of the significance threshold of α = 10^−5^, default parameters were used for spot detection and fitting. To exclude a small number of spurious detections or poorly fit spots, PSFs with background offset value lower than the camera offset level (1,500 counts) were excluded from further analysis; this procedure typically removed approximately 10 % of detected spots.

#### Statistical analysis

A two-sided Wilcoxon rank-sum test (MATLAB function: ranksum) was used to compare distributions with a *p* < 0.05 cutoff for statistical significance.

### *In vivo* lesion tolerance assay

#### Construction of control or lesion-containing plasmids for chromosomal integration

Lesion-containing plasmids were constructed by the gap-duplex method as previously described (Koehl et al., 1989; Pagès et al., 2012). A 13-mer oligonucleotide, 5’-GCAAG<u>TT</u>AACACG-3’, containing no lesion, a TT-CPD or a TT(6-4) lesion (underlined) was inserted into a gapped-duplex pGP1/2 leading to an in frame *lacZ* gene. A 13-mer oligonucleotide, 5’-GAAGACCT<u>G</u>CAGG-3’, containing no lesion or a G-BaP lesion ((-)-trans-anti-benzo[a]pyrene-N^2^-dG)(underlined) was inserted into a gapped-duplex pVP143/pGP9 leading to an in frame *lacZ* gene. Since the G-AAF lesion can be replicated past by two distinct TLS pathways, error free TLS pathway (TLS0) and mutagenic(−2) frameshift TLS pathway (TLS-2) (Figure 5D) (Napolitano et al., 2000), two plasmids were constructed to monitor either TLS events. A 15-mer oligonucleotide containing no lesions or a single G-AAF adduct (underlined) in the *NarI* site (ATCACCGGC<u>G</u>CCACA) was inserted into a gapped-duplex pVP141/142 or pVP143/144, leading to an in frame *lacZ* gene or a +2 frameshift *lacZ* respectively. Therefore, the construct pVP141/142 Nar3^AAF^/Nar+3 monitors TLS0 events, while pVP143/144 Nar3^AAF^/Nar+3 monitors TLS-2 events.

#### Preparation of competent cells for *in vivo* lesion tolerance assay

LB containing kanamycin or chloramphenicol to maintain plasmid pVP135 or pKN13, respectively, both of which express *int-xis* operon (Figure S5A), and 200 µM IPTG are inoculated with 500 ul of an overnight culture. When the culture reached OD_600_ ∼ 0.5, cells were washed in distilled deionized water twice, once in 10% glycerol and finally resuspended in 200 µl of 10% glycerol and frozen in 40 αl aliquots. Lesion-containing plasmids were introduced into these cells by electroporation.

#### Integration protocol – Measurement of lesion Tolerance pathways: TLS and DA

40 µl of electrocompetent cells were transformed with 1 ng of the lesion-containing or lesion-free control plasmids together with 1 ng of pVP146 (transformation control) by electroporation (GenePulser Xcell(tm) from BioRad, 2.5kV – 25µF – 200Ω). Electroporated cells were recovered in 1 ml of SOC medium containing 200 µM IPTG and incubated at 37°C for 1 hour. To measure the transformation efficiency with pVP146 (*tet*^*R*^), a portion of the culture was plated on LB-Agar containing 10 µg/ml tetracycline, and the rest of the culture was plated on LB-Agar containing 50 µg/ml ampicillin and 80 µg/ml X-Gal to select for integrants (Amp^R^) and TLS events (blue-sectored colonies) (Figure S5A). Cells were diluted and plated using an automatic serial diluter and plater (EasySpiral Dilute, Interscience). The integration rates were about 2,000 clones per picogram of plasmid for our parental strain.

To determine TLS and DA at the G-BaP, TT-CPD and TT(6-4) lesions for a given strain, we integrated corresponding heteroduplex plasmids (see above). Blue-sectored colonies represent TLS events, whereas white colonies represent DA. For these lesions, the overall survival was the sum of TLS0 and DA. Each experiment was repeated 3 to 5 times with at least two different batches of competent cells.

In order to determine TLS0, TLS-2 and DA at the G-AAF lesion for a given strain, four integration experiments were performed with the following heteroduplex plasmids: pVP141/142Nar0/Nar+3, pVP141/142Nar3^AAF^/Nar+3, pVP143/144Nar0/Nar+3 and pVP143/144Nar3^AAF^/Nar+3. Overall survival was the sum of TLS0, TLS-2 and DA. Following the integration of pVP141/142Nar3^AAF^/Nar+3, blue-sectored colonies represent TLS0 events, whereas white colonies represent the sum of TLS-2 and DA. Similarly, upon integration of pVP143/144Nar0/Nar+3, blue-sectored colonies represent TLS-2 events, whereas white colonies represent the sum of TLS0 and DA. The integration efficiencies of lesion-containing plasmids were comparable to those of the respective lesion-free control plasmids. Overall survival normalized with the transformation efficiency of pVP146 plasmid, which was added in the same transformation as an internal control, gives the overall rate of lesion tolerance. DA was determined by subtracting TLS0 and TLS-2 from the overall lesion tolerance rate. Each experiment was repeated 3 to 5 times with at least two different batches of competent cells.

### Measuring mutation rates

#### Spontaneous mutation rates – acquisition of rifampicin-resistance

To determine spontaneous mutation rates of various *dnaQ* strains by fluctuation analysis, acquisition of resistance to rifampicin was measured. Single colonies were picked from a fresh streak on LB agar and grown overnight in LB at 37°C with aeration; for each strain, 10 independent colonies were picked and separately cultured. After overnight culture (16 hour), 50 μl of cell suspensions were withdrawn from each culture and plated on a LB agar plate supplemented with 50 μg/ml rifampicin. To determine the number of viable cells, the same cultures were also serially diluted and plated on LB agar. All plates were incubated at 37°C for 24 h before counting colonies. The mutation frequency was calculated as median for 10 cultures divided by a mean value of viable cell count.

#### UV-induced mutation rates – acquisition of rifampicin resistance

An exponentially growing cell culture (OD_700_ ∼ 0.5) was re-suspended in MgSO_4_ (10 mM) and irradiated with total amount of 30 J/m^2^ of UV (254 nm). Immediately following UV irradiation, the irradiated cell suspensions were serially diluted and plated on LB agar for the determination of viable cells. The same irradiated cell suspensions were diluted 20-fold in fresh LB medium and cultured at 37°C with aeration. After culturing for 6h, undiluted cultures were plated on LB agar plates supplemented with 50μg/ml of rifampicin to determine the number of UV-induced *Rif*^*R*^ mutants. To determine the number of viable cells, the same culture were also serially diluted and plated on LB agar plates. All plates were incubated at 37°C for 24 h before counting colonies. The mutation frequency is expressed as the ratio of *Rif*^*R*^ colonies to number of viable cells.

